# Ancient genomes from the last three millennia support multiple human dispersals into Wallacea

**DOI:** 10.1101/2021.11.05.467435

**Authors:** Sandra Oliveira, Kathrin Nägele, Selina Carlhoff, Irina Pugach, Toetik Koesbardiati, Alexander Hübner, Matthias Meyer, Adhi Agus Oktaviana, Masami Takenaka, Chiaki Katagiri, Delta Bayu Murti, Rizky Sugianto Putri, Mahirta, Thomas Higham, Charles F. W. Higham, Sue O’Connor, Stuart Hawkins, Rebecca Kinaston, Peter Bellwood, Rintaro Ono, Adam Powell, Johannes Krause, Cosimo Posth, Mark Stoneking

**Author notes:** These authors contributed equally (Sandra Oliveira, Kathrin Nägele, Selina Carlhoff, Johannes Krause, Cosimo Posth, Mark Stoneking).

## Abstract

Previous research indicates that the human genetic diversity found in Wallacea - islands in present-day Eastern Indonesia and Timor-Leste that were never part of the Sunda or Sahul continental shelves - has been shaped by complex interactions between migrating Austronesian farmers and indigenous hunter-gatherer communities. Here, we provide new insights into this region’s demographic history based on genome-wide data from 16 ancient individuals (2600-250 yrs BP) from islands of the North Moluccas, Sulawesi, and East Nusa Tenggara. While the ancestry of individuals from the northern islands fit earlier views of contact between groups related to the Austronesian expansion and the first colonization of Sahul, the ancestry of individuals from the southern islands revealed additional contributions from Mainland Southeast Asia, which seems to predate the Austronesian admixture in the region. Admixture time estimates for the oldest individuals of Wallacea are closer to archaeological estimates for the Austronesian arrival into the region than are admixture time estimates for present-day groups. The decreasing trend in admixture times exhibited by younger individuals supports a scenario of multiple or continuous admixture involving Papuan- and Asian-related groups. Our results clarify previously debated times of admixture and suggest that the Neolithic dispersals into Island Southeast Asia are associated with the spread of multiple genetic ancestries.

---

Wallacea (Figure 1), a region of deep-sea islands located between the Sunda and Sahul continental shelves^1^, has been both a bridge and a barrier for humans migrating from Asia to New Guinea, Australia, and the Pacific Islands. Modern humans presumably first crossed the Wallacean island chains before reaching Sahul, for which the earliest proposed date is ~65 kyr BP, associated with the Madjedbebe rock shelter in northern Australia^2^. However, this date has been questioned^3^ and the earliest unequivocal dates for modern humans in Sahul are around 47 kyr BP^4-6^. In Wallacea itself, the archaeological record indicates occupation by AMH starting ~46 kyr BP in the southern Wallacean islands^7-9^, ~45.5 kyr BP in Sulawesi^10^, and ~36 kyr BP in the northern Wallacean islands (North Moluccas)^11^.

**Figure 1.**
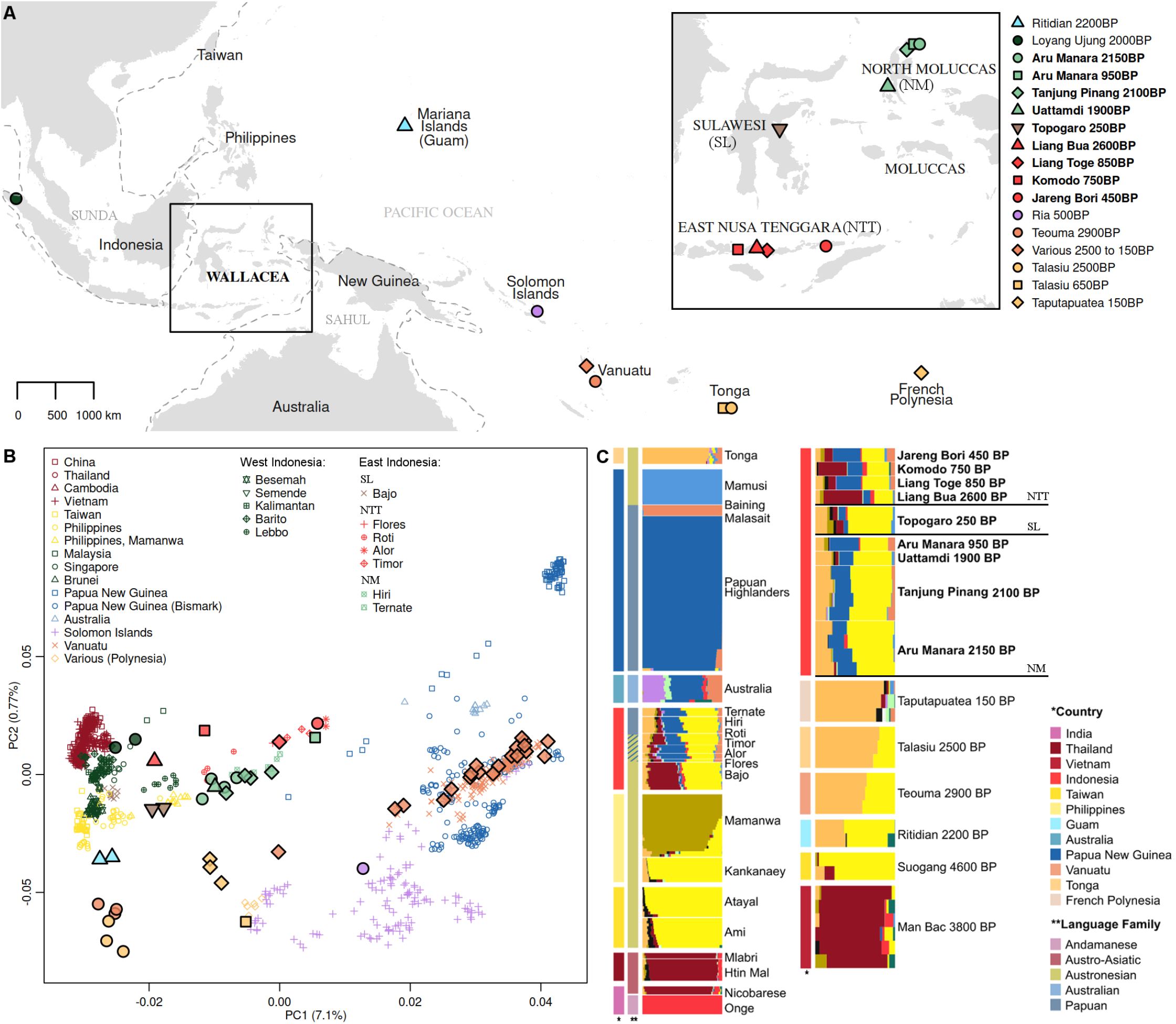
Sample provenience, PC and DyStruct analyses. A) Map showing the location of ancient individuals. B) PCA of publicly available whole genome data merged with Human Origins and Affymetrix 6.0 genotype data. Ancient individuals (shown with a black contour) are projected and their fill color matches the color of present-day individuals from the same geographic area. C) DyStruct results for the same merged dataset as B) displaying only a subset of the individuals included in the full analysis (Figure S2C). Newly generated individuals are highlighted in bold in the legend. Country and Language information are displayed as color bars to the left of the inferred ancestry components.

After a long period of occupation by hunter-gatherer communities, the region was impacted by the Austronesian expansion. Equipped with novel sailing and farming technologies, Austronesian-speaking groups likely expanded out of Taiwan ~4000-5000 yr BP^12-14^ and eventually settled in Island Southeast Asia, Oceania, and Madagascar. Their arrival is generally linked to the earliest appearance of pottery, which dates to ~3500 yr BP in Wallacea^11,15-18^.

During the late Neolithic period and the early Metal Age in Wallacea (2300-2000 yr BP) the maritime trade network intensified further, with a movement of spices such as Northern Moluccan cloves (*Syzygium aromaticum*), bronze drums, and glass beads connecting Wallacea to India and Mainland Southeast Asia^11,17,19-24^.

The contact between Austronesian-speaking groups and the previously established hunter-gatherer communities is still reflected in the linguistic and biological diversity of Wallacea today. Austronesian languages of the Malayo-Polynesian (MP) subgroup are widespread throughout the region^25^, but a few dozen non-Austronesian (i.e., Papuan) languages are also spoken in Wallacea, including in the North Moluccas, Timor, Alor, and Pantar^26^. Additionally, some Austronesian languages spoken in Wallacea show linguistic features acquired from Papuan languages^27^.

Analyses of the genomic composition of present-day groups from the region have shown signals of admixture between Papuan-related ancestry and an Asian-related ancestry most similar to that of present-day Austronesian-speaking groups^28-30^. This dual ancestry is geographically distributed as a gradient of increasing Papuan-related ancestry from west to east^28,30^. Previous studies have estimated the time of admixture of the two ancestries using data from present-day groups^28-30^, providing the first inferences on the direction and rate of spread of genetic ancestry across the islands^28^. However, the time estimates from different studies show discrepancies of more than 3000 yr (Table S1) that cannot be solely attributed to ascertainment biases in the sampled individuals and analyzed markers but are also affected by the methods used to infer the date of admixture^28,29^. In fact, current methods cannot accurately infer admixture dates for complex scenarios involving continuous gene flow or repeated admixture from closely related sources^31^, which could have occurred in the area. As a result, even relatively recent (<4000 yr) demographic movements across Wallacea are not well understood. Resolving the discrepancies between admixture dates has important implications for understanding interactions between Austronesians and indigenous pre-Austronesian populations. Admixture dates close to the archaeological dates proposed for the Austronesian arrival would indicate that admixture occurred soon after contact. In contrast, very recent dates would imply that communities co-existed for a long time before genetically mixing or were mixing for a prolonged time period. Moreover, admixture dates predating the Austronesian arrival would suggest alternative explanations, such as genetic influences from other Asian-related groups in earlier periods^32^.

In this study, we leverage the power of ancient DNA to investigate spatio-temporal patterns of variation within Wallacea during the last ~2500 yr. We provide insights into the time of arrival of the Austronesian-related ancestry, the temporal span of admixture, and the relationship between the ancestry of incomers and the ancestry of other modern human groups from Island Southeast Asia and Oceania. Additionally, we explore the impact and timing of an additional migration from Mainland Southeast Asia to Wallacea.

## Results

We extracted DNA from skeletal remains from 16 individuals dated to ~2600-250 yr BP from eight archaeological sites spanning the North Moluccas, Sulawesi, and East Nusa Tenggara (Figure 1A, Table 1, Table S2). Sequencing libraries were then constructed and capture-enriched for ~1.2 million genome-wide single nucleotide polymorphisms (SNPs)^33^ and the complete mitochondrial genome (mtDNA). The authenticity of ancient DNA was confirmed based on the elevated amounts of deaminated positions at the ends of reads (up to ~45% in non-UDG treated samples) and the short average fragments size (~55 base pairs) (Table S2). Contamination estimates were low (~0.01% for males based on X-chromosomes and ~2% based on mtDNA) (Table S2). The mean mtDNA coverage ranged from ~8X to ~900X (Table S2), while the number of SNPs obtained for each individual ranged from ~123,000 to ~1 million (Table 1).

**Table 1.** Ancient samples from Wallacea included in this study. MtDNA and Y-chromosome haplogroups are colored according to their most likely origin based on previous publications: red – Asian, blue – Australo-Papuan.

| Sample name | Island, region | C14 date ± SD (yrs BP) | Assigned group | Sex | mtDNA haplogroup | Y chromosome haplogroup | Number of SNPs |
| --- | --- | --- | --- | --- | --- | --- | --- |
| AMA001 | Morotai, North Moluccas | 2258 ± 30 | Aru Manara 2150 BP | M | **B4a1a1** | **C1b1a2b** | 255,458 |
| AMA003008 | Morotai, North Moluccas | 2130 ± 24 | | F | **Q1d** | | 576,009 |
| AMA004 | Morotai, North Moluccas | 2009 ± 24 | | F | **M73a** | | 933,715 |
| AMA005 | Morotai, North Moluccas |  |  | F | **B4a1a1** |  | 209,933 |
| AMA009 | Morotai, North Moluccas | 968 ± 20 | Aru Manara 950 BP | F | **Q1d** | | 935,157 |
| TanjungPinang1 | Morotai, North Moluccas | 2090 ± 180 | Tanjung Pinang 2100 BP | M | **Q** | **S1d1~** | 870,652 |
| TanjungPinang2 | Morotai, North Moluccas |  |  | M | **Q1** | **O2a2b2a2b2** | 939,665 |
| TanjungPinang4 | Morotai, North Moluccas |  |  | M | **Q** | **S1a1b1d2b~** | 897,982 |
| TanjungPinang6 | Morotai, North Moluccas |  |  | M | **B4a1a** | **O2a2b2a2b2** | 1,028,190 |
| Uattamdi1 | Kayoa, North Moluccas | 1915 ± 27 | Uattamdi 1900 BP | M | **E1a1a1** | **O1a2a1** | 895,300 |
| TOP002 | Sulawesi, Central Sulawesi | 211 ± 24 | Topogaro 250 BP | M | **E2a** | **O2a2a1a2a2** | 697,028 |
| TOP004 | Sulawesi, Central Sulawesi | 324 ± 24 | | M | **E2a** | **M1a** | 249,209 |
| KMO001 | Komodo, East Nusa Tenggara | 726 ± 19 | Komodo 750 BP | F | **B4a1a1** | | 122,610 |
| LIA001002 | Flores, East Nusa Tenggara | 2588 ± 23 | Liang Bua 2600 BP | F | **M17a** | | 873,614 |
| LIT001 | Flores, East Nusa Tenggara | 861 ± 20 | Liang Toge 850 BP | F | **E1a2** | | 623,960 |
| JAB001 | Pantar, East Nusa Tenggara | 457 ± 19 | Jareng Bori 450 BP | F | **M7b1a2a1** | | 848,849 |

MtDNA and Y-chromosome haplogroups were determined based on the mitochondrial genomes and SNP data, respectively, and are reported, together with their associated origin, in Table 1. The haplogroup composition shows that both Asian- and Australo-Papuan-related ancestries were already present in the North Moluccas ~2150 yrs BP. Furthermore, two North Moluccas individuals dating to ~2150-2100 yrs BP carry mtDNA and Y chromosome haplogroups associated with different ancestries, indicating that admixture started before that period. In comparison to the individuals from East Nusa Tenggara and Sulawesi, those from North Moluccas show a higher proportion of mtDNA lineages that connect them to Near Oceania, as attested by the Q haplogroups characteristic of Northern Sahul ^34^ and by the so-called “Polynesian pre-motif” (B4a1a/B4a1a1)^35^. None of the individuals from Sulawesi or East Nusa Tenggara carry Australo-Papuan-related mtDNA haplogroups, which are found there today^36,37^, likely reflecting the small number of ancient individuals analyzed.

To explore the genome-wide patterns of variation in the ancient individuals from Wallacea, we performed principal component analysis (PCA) based on two different sets of present-day populations from Asia and Oceania (see methods; Figure 1B and Figure S1). The ancient Wallaceans cluster between populations from Papua New Guinea and Asia, together with present-day populations from their respective geographical areas. However, the trajectory outlined by individuals from the northern (North Moluccas) vs. southern (East Nusa Tenggara) islands is slightly different, suggesting they may have distinct genetic histories. Ancient individuals from East Nusa Tenggara cluster on a cline towards mainland Asians and some Western Indonesian groups, while ancient individuals from the North Moluccas align on a trajectory towards present-day Taiwanese and Philippine populations or even towards ancient individuals from Guam 2200 BP, Vanuatu 2900 BP, and Tonga 2500 BP. These ancient individuals were all previously shown to have almost exclusively Austronesian-related ancestry^38,39^.

We next used a model-based clustering method (DyStruct) to infer shared ancestry, considering the dating of the individuals used in the analysis^40^. The results for the best supported number of clusters in each of the tested datasets (Figure S2A-B) show that ancient individuals from Wallacea share ancestry with Papuan-speaking groups from New Guinea (dark blue component) and multiple Asian groups whose ancestry can be partitioned into three main components (Figure 1C; see full results in Figure S2C-D). One component (yellow) is present at high frequencies in Austronesian-speaking groups from Taiwan, Philippines, and Indonesia, as well as ancient individuals from Taiwan; a second component (mango) is maximized in Polynesian-speaking groups from Near Oceania, Tahiti, Tonga, and Samoa, and ancient individuals from the same region; and a third component (dark red) is widespread in present-day and ancient individuals from Southeast Asia. The most striking difference among the ancient Wallaceans is the presence of the Southeast Asian component (dark red) in the ancient East Nusa Tenggara and Sulawesi individuals but the absence of this component in the North Moluccan individuals. A more subtle difference occurs in the relative proportion of the two Austronesian-related components (Figure S3): ancient individuals from Sulawesi and East Nusa Tenggara have a higher relative proportion of the Austronesian-related (yellow) component that predominates in the likely source of the expansion (Taiwan), compared to the ancient individuals from the North Moluccas, who are more similar to groups from the Pacific.

To directly compare the sharing of alleles between ancient individuals from Wallacea and different Asian-related groups, we used *f*-statistics^41^. First, we computed an *f_4_*-statistic of the form *f_4_*(*Mbuti, ancient Wallacean; Amis, test*), where the test group includes the ancient and present-day groups from mainland Asia, Island Southeast Asia, and the Pacific that have no discernible Papuan-related ancestry (Figure S4). A significant positive result indicates that the ancient Wallacean shares more drift with the test group than with the Taiwanese Amis, while a significant negative result indicates that the ancient Wallacean shares more drift with the Amis than with the test group. Our results show that ancient individuals from the North Moluccas share more drift with ancient individuals from Vanuatu (2900 BP) and Tonga (2500BP) than with Amis. In contrast, ancient individuals from Sulawesi and East Nusa Tenggara do not share additional drift with any tested groups. Nonetheless, the higher number of *f_4_*-statistics consistent with zero in tests involving ancient individuals from East Nusa Tenggara (Komodo and Liang Bua) indicates that they share as much drift with Amis as with several other groups, not only from Taiwan and Philippines but also from Southeast Asia or Western Indonesia. This result, together with the identification of an additional ancestry component related to Southeast Asia (Figure S2C-D) in the ancient East Nusa Tenggara and Sulawesi individuals, supports a more complex admixture history in these parts of Wallacea.

We next created a series of two-dimensional plots comparing pairs of *f_4_*-statistics designed to capture any differences between the North Moluccas and East Nusa Tenggara individuals that might reflect differential Asian-related ancestries. All pairs had the form *f_4_*(*Mbuti, test; New Guinea Highlanders, ancient Wallacean*) and always included Amis or ancient Vanuatu (2900 BP) vs. other Asian-related groups as *test*. Since individuals from the North Moluccas lack the Southeast Asian component in the DyStruct analysis, their differences to East Nusa Tenggara individuals regarding the attraction to the Asian-related groups should be maximized by the best proxies for the Southeast Asian ancestry in East Nusa Tenggara. By comparing pairs of present-day groups (Figure S5) and pairs of ancient groups (Figure S6), we conclude that the groups maximizing differences between ancient individuals from Wallacea are the present-day Mlabri or Nicobarese and ancient individuals from Vietnam (Nam Tun 2600BP, Man Bac 3800BP, Mai Da Dieu 4000BP), Cambodia (Vat Komnou 1800BP), Thailand (Ban Chiang 2500BP and 3350BP), Malaysia (Gua Cha Cave 2300BP), and Laos (Tam Pa Ping 3000BP) (Figure 2). These are thus the best proxies among the tested groups and individuals for the Southeast Asian ancestry in East Nusa Tenggara.

**Figure 2.**
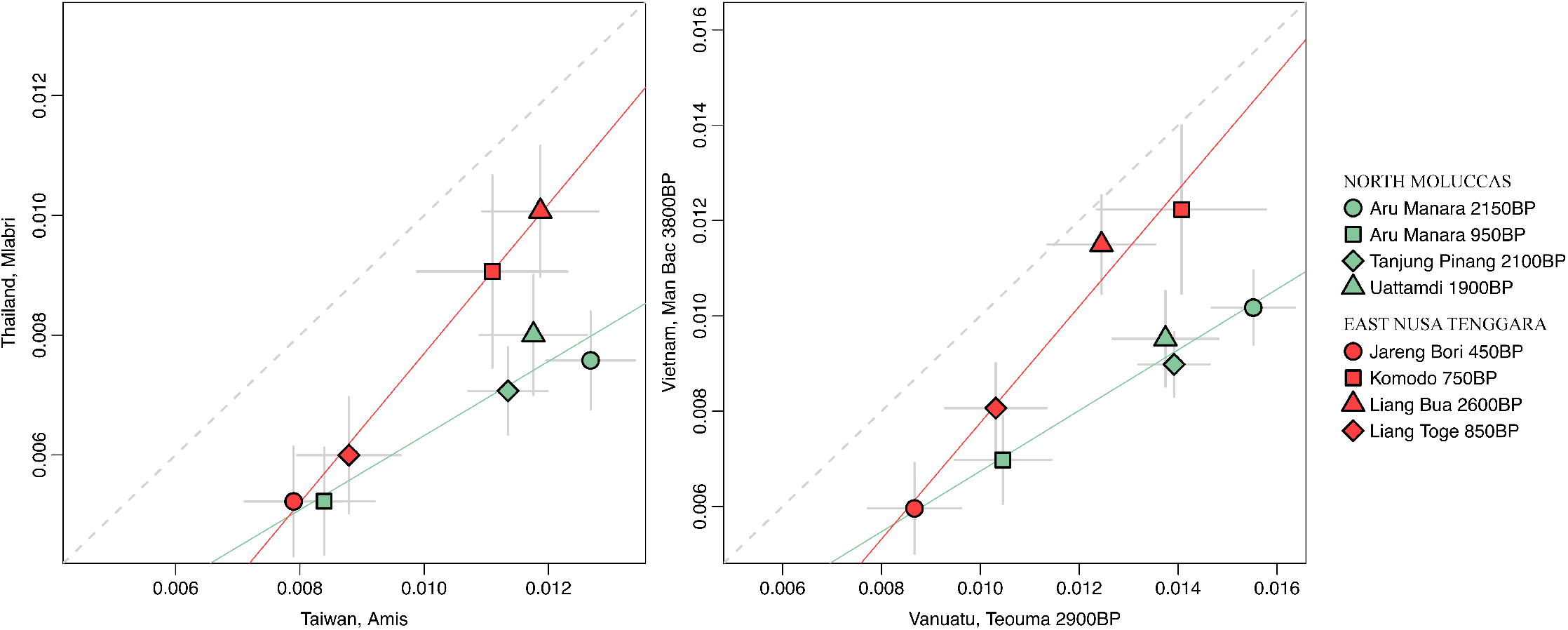
Biplots showing the results of two pairs of *f_4_*-statistics of the form: *f_4_*(*Mbuti, test; New Guinea Highlanders, ancient Wallacea*). The test groups are shown on the x-y axis label. Grey bars show two standard errors in each direction. Linear regression lines for the North Moluccas and East Nusa Tenggara individuals are shown in green and red, respectively. The results for all tested pairs are shown in Figures S5 and S6.

We also investigated the relationships between the newly presented ancient Wallaceans and groups associated with the first colonization of Sahul or Wallacea using an *f*-statistic of the form *f_4_*(*Mbuti, new ancient Wallacean; New Guinea Highlanders, test*). Since the tested groups are present-day Australo-Papuans with no discernable Asian ancestry and a recently published pre-Neolithic individual from Sulawesi (Leang Panninge)^42^, values consistent with zero indicate that ancient Wallaceans are equally related to New Guinea Highlanders and the tested groups, while deviations from zero indicate differences in affinity to the tested group. Most ancient Wallacean individuals show a significantly closer affinity to New Guinea Highlanders than to Australians, the group representing the Bismarck archipelago, or the Leang Panninge individual (Figure S7). The low number of SNPs available for some *f_4_* tests (e.g., tests involving the Leang Panninge individual) and the low amount of Papuan ancestry carried by the Liang Bua and Topogaro individuals likely account for the non-significant differences in affinity with New Guinea Highlanders vs. Bismarcks or Leang Panninge by broadening the confidence intervals (Figure S7) and reducing the power to distinguish between potential sources of this ancestry (Figure S8), respectively. Nonetheless, tests involving the Leang Panninge individual consistently exhibit the lowest *f_4_* values. Therefore, despite being from Wallacea, this ancient individual is a worse proxy for the Papuan-related ancestry of the newly reported ancient Wallaceans than any other tested group.

We further investigated potential differences in ancestry among ancient Wallaceans by modeling ancestry sources and proportions with the qpAdm software^41^. Our results indicate that whereas the ancient individuals from the North Moluccas can be modeled as having both Papuan- and Austronesian-related ancestry, ancient individuals from East Nusa Tenggara and Sulawesi are either consistent with or require a three-wave model, with additional Southeast Asian-related ancestry (Figure 3; Table S3). Despite the cases for which we identify more than one fitting model (*P* > 0.01), the estimated proportions under the model with the highest *P*-value correlate with the proportions of Austronesian, Papuan, and Southeast Asian ancestry inferred by DyStruct (mantel statistic r = 0.97, *P* < 0.001). The ancient East Nusa Tenggara individuals display more inter-island variance in their Papuan- and Southeast Asian-related ancestries (s^2^ = 0.026 and 0.046, respectively) compared to their Austronesian-related ancestry (s^2^ = 0.003).

**Figure 3.**
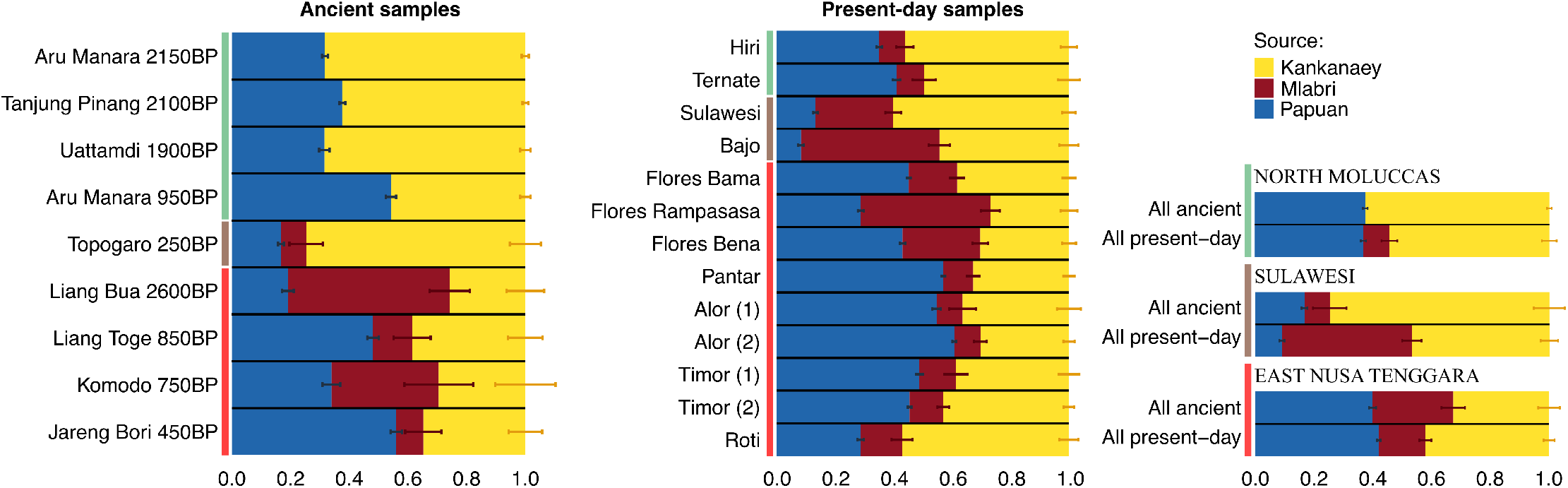
Ancestry proportions for the model with the highest *P*-value in each group. Individuals from the North Moluccas, Sulawesi, and East Nusa Tenggara are marked with green, brown, and red vertical bars, respectively. Horizontal bars show standard errors. The number following the name of the present-day individuals indicates the genotyping array used: (1) Affymetrix 6.0 and (2) Affymetrix Axiom Genome-Wide Human.

A comparison between the ancestry composition of ancient and present-day individuals from the same region (Figure S3; Table S3) suggest that a small part (~8%) of the Austronesian-related ancestry of the ancient North Moluccas individuals was replaced by Southeast Asian ancestry in present-day groups, masking former differences between regions of Wallacea. The present-day groups from Sulawesi and East Nusa Tenggara can be modeled by the same three ancestry components found in the ancient individuals from those regions. However, the ancestry proportions of ancient and present-day groups show some differences at the regional level, which could be explained by ancestry shifts over time, or could also reflect the small sample sizes.

To gain insights into the relative order of admixture events between different ancestries through time in the northern and southern Wallacean islands, we used an approach called Admixture History Graph (AHG)^43^ that relies on differences in the covariance between the ancestry components inferred by the DyStruct analysis (Table S4-6). The AHG, applied to both ancient and present-day data from East Nusa Tenggara, suggests that the admixture of Southeast Asian- and Papuan-related ancestries occurred before the arrival of the Austronesian-related ancestry (Table S6). An analogous test based on the three main ancestry components observed among individuals from the North Moluccas (one Papuan and two Austronesian-related) does not provide compelling evidence of backflow from Austronesian-related groups of the Pacific, as the AHG infers that Papuan ancestry was introduced into a population that already had the two Austronesian-related components (Table S6). This result suggests that drift had a more significant role in the occurrence and distribution of the two Austronesian-related components.

Finally, we investigated the timing of admixture using the software DATES (Figure S9), which can be applied to ancient DNA data from single individuals and has been shown to return estimates that are as accurate as those estimated by other methods based on multiple individuals^44^. Since we estimate admixture dates for individuals from different periods, we expect to gain more insights about admixture through time and reconcile the time estimates from different methods, despite the challenges associated with dating complex scenarios. Using Papuans and a pool of Asian groups as sources, we found that estimates for the oldest individuals from the Northern Moluccas (2150 BP) and East Nusa Tenggara (2600 BP) are very similar (~3000 years BP, adjusting for the archaeological age of each sample; Figure 4), approaching archaeological dates for the arrival of the Austronesians in Wallacea. However, ancient individuals from more recent periods display more recent estimates. Contrary to present-day groups from East Nusa Tenggara, present-day groups from the North Moluccas show even younger admixture dates (~1400 BP) than the ancient individuals from the same region (Figure 4). This pattern probably reflects differential admixture pulses (and/or continuous gene flow) since the initial contact between Austronesians and Papuan-related groups, as suggested by the changes in ancestry composition or ancestry proportions over time (Figure 3).

**Figure 4.**
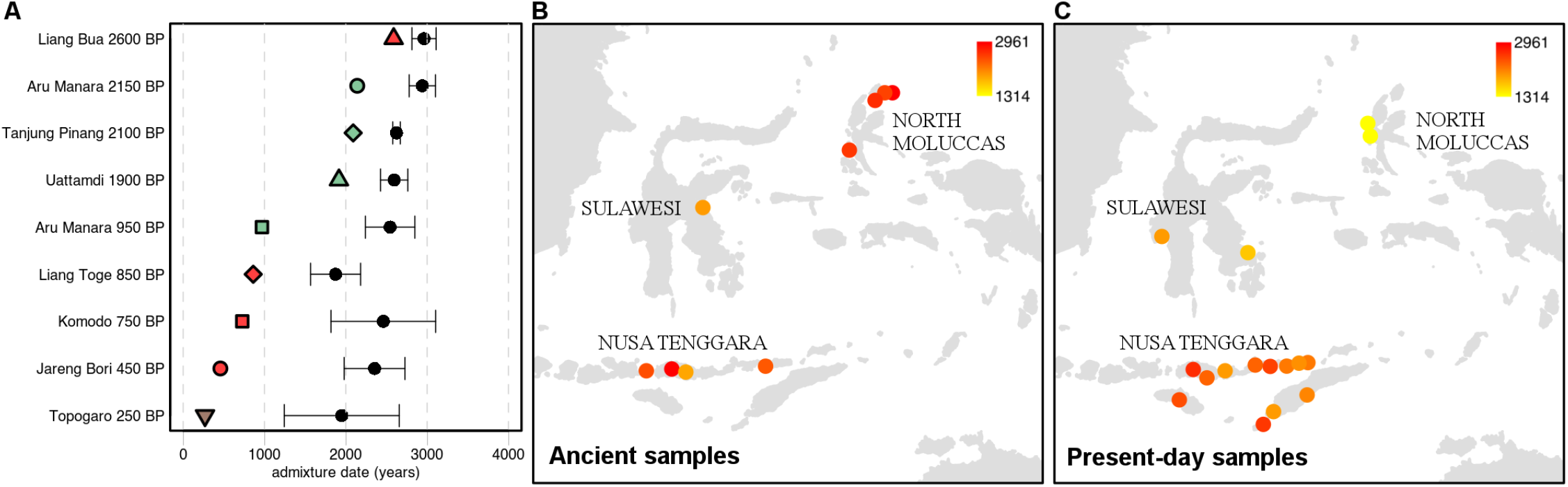
Admixture estimates. A) Admixture date point estimates and standard errors are shown in black. The individuals age is indicated with filled symbols: green – North Moluccas, brown – Sulawesi, red – East Nusa Tenggara. B-C) Spatial distribution of admixture date estimates for ancient and present-day individuals, with admixture dates depicted according to the heat plot.

## Discussion

This study greatly increases the amount of ancient genomic data from Island Southeast Asia, a tropical region unsuitable for DNA preservation but particularly important for understanding human population interactions and admixture. Notably, we report the first genomic data from an ancient human from the Liang Bua cave on the island of Flores, the same site that provided the only known remains of *Homo floresiensis*. Although the amount of genomic data recovered precluded detailed investigation into admixture with archaic humans, the fact that authentic ancient DNA could be recovered from 2600 yrs old remains from Liang Bua suggests that such investigations could be possible for other ancient human remains from this site.

Our results reveal striking regional variation in the ancestry of Wallacean individuals and provide insights into the temporal window and nature of events that shaped them. Some of the variation is found among the Austronesian-related ancestry of ancient northern vs. southern Wallaceans. The most remarkable differences are associated with ancestry contributions from Mainland Southeast Asia that were probably already part of the East Nusa Tenggara genomic landscape when Austronesians arrived. The Australo-Papuan ancestry of Wallaceans, on the other hand, supports some shared or parallel histories.

### Australo-Papuan ancestry in Wallacea

Based on our analysis, all newly presented ancient Wallacean individuals are genetically closer to present-day Australo-Papuans than to the recently published pre-Neolithic Leang Panninge individual (~7 kyr BP) from Sulawesi^42^, suggesting little direct continuity between pre-Neolithic and post-Austronesian Wallaceans. Additionally, the Australo-Papuan ancestry of the newly presented Wallaceans is closer to the ancestry of present-day Papua New Guineans than indigenous Australians. This proximity suggests that either the group that gave rise to Australians split first, or there was contact between Wallacea and Papua New Guinea after their initial settlement. The second scenario is supported by a recent mtDNA study which inferred two major influxes of Papuan ancestry into Wallacea, one following the Last Glacial Maximum ~15 kyr, the other associated with Austronesian contact ~3 kyr^45^.

Previous studies have also reported elevated amounts of Denisovan ancestry in present-day Wallaceans, which correlate with the amount of Papuan-related ancestry^46^. We confirmed that the same relationship holds for the ancient Wallaceans included in this study by plotting estimated Papuan vs. Denisovan ancestry (Figure S10), thus supporting the idea that their Denisovan-related ancestry was likely contributed via Papuan-related admixture.

### Early Southeast Asian ancestry in East Nusa Tenggara

The area to the east of the Wallace line is generally assumed to have been mostly isolated and shaped by two main streams of ancestry: one related to the first settlement of Sahul and another associated with the Austronesian expansion^28-30^. The results presented here show that the genetic variation of ancient individuals from East Nusa Tenggara also requires ancestry contributions from Mainland Southeast Asia. The Admixture History Graph (AHG) analysis further suggests that this ancestry likely admixed with Papuan-related ancestry before the arrival of Austronesian-related ancestry. The inferred order of events makes it unlikely that the Southeast Asian and Austronesian-related ancestries were introduced together in the south of Wallacea from Western Indonesia, where both ancestries are found. Instead, it seems that human groups from Mainland Southeast Asia crossed into southern Wallacea before the Austronesian-related groups spread into the region.

The broad geographical distribution of ancient and present-day groups best matching the Mainland Southeast Asian ancestry found in the south of Wallacea raises questions about the actual origin(s) of peoples that reached those islands. The best present-day proxies, the Mlabri from Thailand/Laos and the Nicobarese from the Nicobar islands, speak Austroasiatic languages and are genetically similar, with the Nicobarese displaying some additional ancestry related to non-Austroasiatic Onge (from the neighboring Andaman Islands)^47^. Even though the Mlabri are a hunter-gatherer group, their ethnogenesis has been attributed to a strong founder event from an ancestral agricultural population^48^. The fact that both the Mlabri and Nicobarese have been relatively isolated compared to other Southeast Asian groups that recently experienced extensive admixture^49,50^, might explain why they are the best present-day proxies for the Mainland Southeast Asian ancestry in southern Wallacea, without being necessarily connected to the inferred migration event. Moreover, ancient individuals from all across Mainland Southeast Asia are also equivalent proxies for this ancestry in southern Wallacea (Figure 2 and S6); thus, there is no clear link between any specific ancient group from Southeast Asia and the actual source that contributed ancestry to East Nusa Tenggara.

The only documented languages in East Nusa Tenggara belong to the Austronesian family (Malayo-Polynesian subgroup) or the Papuan language cluster, and no significant influences from Mainland Southeast Asian language families have been reported in southern Wallacea. While all populations in Island Southeast Asia to the west of Wallacea currently speak Austronesian languages, an Austroasiatic influence has been suggested in some Borneo languages^51^, and Borneo individuals carry a relatively high degree of Mainland Southeast Asian genetic ancestry (Figure S2). However, no linguists have suggested a prior presence of Austroasiatic languages in East Nusa Tenggara, and neither are there direct archaeological traces of pre-Austronesian contact between this region and Mainland Southeast Asia.

The earliest evidence of interaction likely comes from the appearance of Dong Son bronze drums, which spread to southern but not northern Wallacea around the early centuries AD, following early maritime trade routes^52^. These drums probably originated in northern Vietnam or adjacent provinces of southern China^52^. Although we cannot rule out some Southeast Asian ancestry contributions from the Dong Son period (or even later) for the younger East Nusa Tenggara individuals, the high amount of Southeast Asian ancestry in the Liang Bua individual (2600 BP) and our AHG inferences support its earlier presence in southern Wallacea. Ancient DNA from older time periods will help to clarify the time of arrival of this ancestry.

### The relationship between the Austronesian ancestry of the North Moluccas and the Pacific

The fine-scale structure observed among Austronesian-related groups from Island Southeast Asia and the Pacific, and the higher genetic proximity of the ancient North Moluccas individuals to the latter, are pertinent for earlier considerations of the role of the North Moluccas in dispersals to the Pacific^20^. When analyzed through sophisticated seafaring and climatic models, the North Moluccas appears to be one of the most likely starting points for settlers that ventured into the Palau or Mariana Islands (Western Micronesia)^53,54^. Their geographical setting also led archeologists to search the region for pottery that could be considered ancestral to the Lapita cultural complex (distributed from the Bismarck archipelago to New Caledonia in the south and Samoa in the east), as well as the Marianas Redware culture^11,18^. However, to date no evidence connects the North Moluccas red-slipped pottery to either of these material cultures, and ancient DNA from Guam supports an origin for the settlement of the Mariana Islands from the Philippines^39^. Overall, the North Moluccas red-slipped Neolithic pottery is closer to that of the Talaud Islands, Northern and Western Sulawesi, North Luzon, Batanes, and Southeastern Taiwan^11^.

Our genetic results are compatible with a route through the North Moluccas, leading to the settlement of the Mariana Islands and the Lapita-related dispersals. Under a relatively simple expansion scenario, without significant back migration, the increasing amounts of Austronesian ancestry characteristic of the Pacific (and decrease of ancestry characteristic of Taiwan/Philippines) from the ancient North Moluccas to Guam (~2200 BP), Vanuatu (~2900 BP), and Tonga (~2500 BP) could reflect their relative position along the peopling wave that eventually reached the Eastern parts of the Pacific (Figure S3). Yet, the position here refers to the split order of groups, without any necessary attachment to their geographical location. Therefore, it is possible that the higher proximity between the North Moluccas and groups from the Pacific, compared to East Nusa Tenggara or Sulawesi, simply reflects their more recent ancestry tracing back to a common Austronesian source, regardless of its location. This scenario also implies that the Austronesian-related ancestry found in East Nusa Tenggara or Central Sulawesi is somewhat differentiated from that found in the North Moluccas. Furthermore, we cannot exclude the possibility of more complex migration scenarios (e.g., involving back-migrations).

It is also important to consider that the dates of the oldest individuals from the North Moluccas (2150 BP in Morotai and 1900 BP in Kayoa island) overlap with the start of the Early Metal age ~2300-2000 yr BP in this region^11^. This period is characterized by the appearance of copper, bronze, and iron artifacts and glass beads in the region, as well as the spread of pottery into Morotai island. Thus, these individuals might not be good representatives of the first wave of Austronesians, thought to have reached Kayoa island 3500 yr BP^11,18^, but instead might feature additional genetic influences brought by later contacts.

The Austronesian (Malayo-Polynesian major subgroup) languages of the Northern Moluccas are part of the South Halmahera-West New Guinea (SHWNG) regional subgroup. According to Blust^55^, the SHWNG languages are closer to Oceanic languages - the most clearly defined MP branch - than to any other Western MP subgroup, have a time depth of ~2000 yr, and might represent a replacement movement from the East^55,56^. Additionally, the language phylogeny of Gray et al.^13^ places the SHWNG languages as the closest outgroup to all Oceanic languages and further estimates their divergence to around 3500 yr BP, whereas the languages spoken in East Nusa Tenggara are an outgroup to both SHWNG and Oceanic languages. Thus, the closer genetic proximity of the ancient North Moluccas to Oceanians, compared to East Nusa Tenggarans, is also mirrored in their linguistic relationships.

### The timing of Admixture

Besides providing direct evidence of Austronesian-Papuan contact prior to 2150 yr BP in the North Moluccas and 2600 yr BP in East Nusa Tenggara, the oldest individuals gave admixture date estimates close to 3000 years BP. This period is slightly younger than the earliest archaeological traces of the Neolithic (Austronesian) arrival in the North Moluccas (~3500 years BP for Kayoa Island^11,18^) but older than the adoption of pottery on Morotai Island (~2300-2000 years BP), where the oldest North Moluccan individuals in this study were found^11,57^. However, it is similar to some of the earliest secure dates from East Nusa Tenggara (~3000 years BP for eastern Flores^17^. Some of the previous studies conducted on present-day eastern Indonesian populations suggested that this admixture lagged about a millennium after the arrival of Austronesian populations^30^. Our admixture analysis for ancient individuals, and the comparison with present-day data, provides an alternative explanation, and helps to clarify previous debates concerning times of admixture^28–30,32,58^. The decreasing trend in admixture estimates from the oldest individuals until present-day populations is a strong indicator that admixture occurred in multiple pulses, or even continuously, over a considerable time period. Therefore, even our oldest estimates might not correspond to the actual start of admixture but rather to a more recent time due to the bias introduced by continuous admixture.

Evidence of interaction between populations over time comes from multiple sources. In the Metal Age, admixture might have been facilitated by emergent maritime networks and spice trade interactions^11^. In the North Moluccas, this period not only corresponds to a more rapid spread of material culture between regions^11,21,22,57^, but also to the inferred period of language leveling or radiation described for both the Austronesian (SHWNG) and Papuan (West Papuan Phylum, Northern Halmahera Stock) languages of the North Moluccas^11^. The historical socio-economic systems of the North Moluccas and western Papua also brought together Papuan-speaking resident populations and a Malay-speaking elite^11^, reinforcing that mixing could have occurred until very recent times, and have involved diverse peoples. In contrast to the North Moluccas, the East Nusa Tenggara and Sulawesi ancient and present-day individuals do not show genetic traces of very recent contact. Still, their demographic history was nonetheless characterized by a long-term process of admixture involving at least two Asian-related ancestries.

These findings supporting ongoing contact in Wallacea not only improve our understanding of the history of the region, but also have important implications for efforts that use present-day genomic data to discern the direction and number of human migrations to the Australo-Papuan region (e.g. ref. ^59^). In particular, failure to take this ongoing contact into consideration may result in wrongly considering the genetic affinity between Papuans and northern vs. southern Wallaceans to reflect ancestral relationships of these groups rather then differences in the degree of contact. Overall, our findings suggest different histories for northern vs. southern Wallaceans that reflect differences in contact with Southeast Asia, in the temporal span of contact with Papua New Guinea, and perhaps even with different Austronesian-related groups. Future ancient DNA studies involving individuals from earlier periods will help to improve our understanding of the demographic changes occurring both before and after the arrival of farming in Wallacea.

## Methods

### Sampling

All samples were processed in dedicated ancient DNA laboratories at the Max Planck Institute for the Science of Human History (Jena) and the Max Planck Institute for Evolutionary Anthropology (Leipzig), Germany. In Jena, the petrous bone of samples AMA001, AMA004, and AMA009 was first drilled from the outside, identifying the position of the densest part by orienting on the internal acoustic meter and drilling parallel to it into the target area to avoid damaging the semicircular ducts^60^ (protocol: dx.doi.org/10.17504/protocols.io.bqd8ms9w). After that, the petrous bone was cut along the margo superior partis petrosae (crista pyramidis) and 50 – 150 mg bone powder were drilled from the densest part around the cochlea^61^. All other elements processed in Jena (AMA003, AMA005, AMA008, JAB001, KMO001, LIA001, LIA002, LIT001, TOP002, TOP004) were sampled by cutting and drilling the densest part. In Leipzig, the Tanjung Pinang and Uattamdi specimens were sampled by targeting the cochlea from outside. For this, a thin layer of surface (~1 mm) was removed with a sterile dentistry drill. Small holes were then drilled into the cleaned areas, yielding between 42 and 63 mg of bone powder. Detailed information on the analysed samples, radiocarbon dating, and archaeological context are provided in the Supplementary Information and in Tables S2 and S7.

### DNA Extraction

DNA extraction in both laboratories was carried out using a silica-based method optimized for the recovery of highly degraded DNA^62^, with modifications described elsewhere^63^ and large-volume silica spin columns and binding buffer ‘D’ for DNA binding. In Jena, the second elution of DNA from the spin column was carried out using a fresh aliquot of elution buffer, for a total of 100 μl DNA extract, whereas in Leipzig the same aliquot of elution buffer was loaded twice, for a total of 50 μl DNA extract (protocol: dx.doi.org/10.17504/protocols.io.baksicwe).

### Library preparation

In Jena double stranded DNA libraries were built from 25 μl of DNA extract in the presence of uracil DNA glycosylase (UDG-half libraries), following a protocol that uses the UDG enzyme to reduce, but not eliminate, the amount of deamination-induced damage towards the ends of aDNA fragments^64^. Negative and positive controls were carried alongside each experiment (extraction and library preparation) (protocol: dx.doi.org/10.17504/protocols.io.bmh6k39e). Libraries were quantified using the IS7 and IS8 primers^65^ in a quantification assay using a DyNAmo SYBP Green qPCR Kit (Thermo Fisher Scientific) on the LightCycler 480 (Roche). Each aDNA library was double indexed^66^ in parallel 100 μl reactions using PfuTurbo DNA Polymerase (Agilent) (protocol: dx.doi.org/10.17504/protocols.io.bakticwn). The indexed products for each library were pooled, purified over MinElute columns (Qiagen), eluted in 50 μl TET and again quantified using the IS5 and IS6 primers^65^ using the quantification method described above. 4 μl of the purified product were amplified in multiple 100 μl reactions using Herculase II Fusion DNA Polymerase (Agilent) following the manufacturer’s specifications with 0.3 μM of the IS5/IS6 primers. After another MinElute purification, the product was quantified using the Agilent 2100 Bioanalyzer DNA 1000 chip. An equimolar pool of all libraries was then prepared for shotgun sequencing on the Illumina HiSeq 4000 platform using a SR75 sequencing kit. Libraries were further amplified with IS5/IS6 primers to reach a concentration of 200-400 ng/μl as measured on a NanoDrop spectrophotometer (Thermo Fisher Scientific). In Leipzig, single-stranded DNA libraries were prepared without UDG treatment using the Bravo NGS workstation B (Agilent Technologies), followed by library quantification, amplification and double-indexing, as described in detail in ref. ^67^.

### Targeted enrichment and high-throughput sequencing

Mitochondrial DNA capture^68^ was performed on screened libraries which, after shotgun sequencing, showed the presence of aDNA, highlighted by the typical C to T and G to A substitution pattern towards the 5’ and 3’ molecule ends, respectively. Furthermore, samples with a percentage of human DNA in shotgun data around 0.1% or greater were enriched for a set of 1,237,207 targeted SNPs across the human genome (1,240K capture) as described previously^33^. The enriched DNA product was sequenced on an Illumina HiSeq 4000 instrument with 75 cycles single-reads or 50 cycles pair-end-reads using the manufacturer’s protocol (in Jena), or on a HiSeq 2500 with 75 paired-end-reads (in Leipzig). The output was demultiplexed using in-house scripts requiring either a perfect match of the expected and observed index sequences (Leipzig samples) or allowing a single mismatch between the expected and observed index sequences (Jena samples).

### Genomic data processing

Pre-processing of the sequenced reads was performed using EAGER v.1.92.55^69^. The resulting reads were clipped to remove residual adaptor sequences using *Clip&Merge*^70^ and *AdapterRemoval* v.2^71^. Clipped sequences were then mapped against the human reference genome hg19 using the Burrows– Wheeler Aligner (BWA) v.0.7.12^72^, disabling seeding (-l 16500) and allowing for 2 mismatches (–n 0.01). Duplicates were removed with DeDup v. 0.12.2^69^. Additionally, a mapping quality filter of 30 was applied using SAMtools v.1.3^73^. Different sequencing runs and libraries from the same individuals were merged, duplicates removed, and sorted again using SAMtools v.1.3^73^. Genotype calling was performed separately for trimmed and untrimmed reads using pileupCaller v.8.6.5 (https://github.com/stschiff/sequenceTools), a tool that randomly draws one allele at each of the targeted SNPs covered at least once. For the UDG-treated libraries produced in Jena, 2 bases were trimmed on both ends of the reads. For libraries produced in Leipzig (without UDG treatment) the damage-plots were inspected to determine the number of bases to trim from each read. For all libraries, the residual damage extended 8bp into the read, after which it was below 0.05%, and trimmed accordingly. We combined the genotypes keeping all transversions from the untrimmed genotypes and transitions only from the trimmed genotypes to eliminate problematic, damage-related transitions overrepresented at the ends of reads. The generated pseudo-haploid calls were merged with previously published ancient data^38,39,42,47,74-78^ present-day genomes from the Simons Genome Diversity Project^79^ and worldwide populations genotyped on the Affymetrix Human Origins array^38,41,49,74,75,80-85^. For PCA and DyStruct analyses we additionally merged the data with populations from Island Southeast Asia genotyped on the Affymetrix 6.0^46,86^ (dataset 1) or the Affymetrix Axiom Genome-Wide Human Array^30^ (dataset 2), filtering out SNPs with a missing rate higher than 10%. Related individuals were excluded if they exhibited a proportion of identity by descent (IBD) higher than 0.3, computed in PLINK v.1.9^87^ as P(IBD=2) + 0.5*P(IBD=1). We additionally pruned datasets 1 and 2 for linkage disequilibrium with PLINK v.1.9, removing SNPs with r^2^ > 0.4 in 200 kb windows, shifted at 25 SNP intervals. After pruning, a total of 89,597 and 65,880 SNPs remained in datasets 1 and 2, respectively.

Y-chromosomal haplogroups were identified by calling the SNPs covered on the Y-chromosome of all male individuals using the pileup from the Rsamtools^88^ package and by analysing the SNPs overlapping with the ISOGG SNP index v.14.07 (https://isogg.org/tree) as described in ref. ^89^.

### Authentication of ancient DNA

The typical features of ancient DNA were inspected with *DamageProfiler* v.0.3.1 (http://bintray.com/apeltzer/EAGER/DamageProfiler)^69^. Sex determination was performed by comparing the coverage on the targeted X-chromosome SNPs to the coverage on the Y-chromosome SNPs, both normalized by the coverage on the autosomal SNPs^70^ (Table S2). For male individuals, ANGSD v.0.919 was run to measure the rate of heterozygosity of polymorphic sites on the X-chromosome after accounting for sequencing errors in the flanking regions^90^. This provides an estimate of nuclear DNA contamination in males, as they are expected to have only one allele at each site. For both male and female individuals, mtDNA-captured data were used to jointly reconstruct the mtDNA consensus sequence and estimate contamination levels with contamMix^68^ (Table S2) using an in-house pipeline (https://github.com/alexhbnr-mitoBench-ancientMT39).

### Statistical analyses

Principal component analyses were carried out using *smartpca* v.10210^91^ based on present-day Asian and Oceanian populations from datasets 1 and 2. Ancient individuals were projected onto the calculated components using the options ‘lsqproject: YES’ and ‘numoutlieriter: 0’. We used DyStruct v.1.1.0^40^ to infer shared genetic ancestry from time-series genotype data. The uncalibrated radiocarbon dates of each ancient sample were converted to generations, assuming a generation time of 29 years^92^. For each dataset (1 and 2), we performed 25 independent runs, using 2 to 15 ancestral populations (K). To compare runs for different values of K, a subset of loci (5%) was held out during training and the conditional log likelihood was subsequently evaluated (Figure S2A-B). Within the best K, the run with the highest objective function was selected (Figure S2C-D).

To formally test population relationships we used *f_4_* statistics, implemented in the ADMIXTOOLS software^41^. This analysis was carried out using the admixr R package^93^. We used *qpWave* v.410^94^ and *qpAdm* v.650^41^ to test two- and three-wave admixture models, following a “rotating” strategy^95^. A reference set of populations was chosen to represent diverse human groups and include potential source populations for the ancient Wallacean individuals: Mbuti, English, Brahui, Onge, Yakut, Oroqen, Lahu, Miao, Dai, Khomu, Denisova, Papuan, Kankanaey, Mlabri. We rejected models if their *P*-values were lower than 0.01, if there were negative admixture proportions, or if the standard errors were larger than the corresponding admixture proportion. When more than one model was accepted (Table S3), the estimated admixture proportions under the model with the highest *P*-value was preferred and used in subsequent analyses (Figure S8 and S10) because the results better matched the DyStruct ancestry proportions, and the ability to reject models might be affected by several factors (e.g., the ancestry proportion, the quality of the target sample, the combination of ancient and present-day samples in the same analysis, etc). The correlation between ancestry proportions inferred with *qpAdm* and DyStruct was assessed using a Mantel test with 10,000 permutations of the distance matrix to determine significance.

The relative order of the mixing of different ancestries was inferred using the Admixture History Graphs (AHG) approach described previously^39^. We used the DyStruct proportions for each ancient and present-day Wallacean individual included in dataset 1 and 2 (Table S4-5) to calculate the covariances between ancestry components as indicated in Table S6. The time since admixture was estimated based on the decay of ancestry covariance using the software DATES v.753^44^, with parameters: binsize: 0.001; maxdis: 1.0; jackknife: YES; qbin: 10; runfit: YES; afffit: YES; lovalfit: 0.45; mincount: 1. To maximize the number of SNPs included in the analysis and have equal sample sizes, we used as sources 16 Papuan individuals and 16 Asian-related individuas (2 Ami, 1 Atayal, 2 Kankanaey, 5 Dai, 2 Dusun, 2 She, 2 Kinh) with data covering the ~1,240,000 SNPs captured in the ancient samples.

## Acknowledgements

We thank Rita Radzeviciute, Antje Wissgott, the MPI-EVA lab technicians, and the MPI-EVA Sequencing and Bioinformatics Groups for their excellent support, Choongwon Jeong for valuable comments, and Leonardo Iasi for helpful discussions on admixture dating. This research was supported by the Max Planck Society. KN, SC and AP were supported by the European Research Council Starting Grant ‘Waves’ (ERC758967). Excavations of the samples from Liang Toge, Liang Bua and Komodo were part of a New Zealand Fast-Start Marsden Grant (18-UOO-135). The research conducted on the Jareng Bori site was part of a joint project between the Australian National University (Canberra, Australia) and Universitas Gadjah Maja (Yogyakarta, Indonesia) funded by ARC Laureate Project FL120100156.

## Author contributions

RO, PB, RK, TK, and SH contributed archaeological material, collected with the critical support of AAO, MT, CK, DBM, RSP, M, and SOC. TH, CFWH, RK, and PB contributed radiocarbon data. KN, SC and MM conducted the ancient DNA laboratory work. SO, KN, SC, IP, and AH performed the genetic analysis. SO and KN wrote the manuscript with input from all authors. MS, CP, and JK concieved and coordinated the study.

