## Supplementary information for "Ancient genomes from the last three millennia support multiple human dispersals into Wallacea"

### *1. Ethics statement*

The human remains analyzed in this study are considered the cultural heritage of the originating countries. Appropriate research permits and export approvals were obtained from Indonesia to conduct the current research. For the skeletal material from Tanjung Pinang and Uattamdi, the research was undertaken as part of a collaborative project between Pusat Penelitian Arkeologi Nasional (Jakarta) and the Australian National University, under Lembaga Ilmu Pengetahuan Indonesia Research Permits 6939/S.K./1990, 307/I/KS/1994, and 10290/V3/KS/1995. For the skeletal material from Aru Manara, the research was undertaken as part of a collaborative project between Pusat Penelitian Arkeologi Nasional (Jakarta) and the Tokai University, under Kementarian Riset dan Teknologi Research Permits 0291/SIP/FRP/VIII/2011 and Pusat Penelitian Arkeologi Nasional export permit UM.001/2595/PAN/KPK/IX/2012, 2634/H5/TU/2017. For the skeletal material from Topogaro, the research was undertaken as part of a collaborative project between Pusat Penelitian Arkeologi Nasional (Jakarta) and the Tokai University, under Kementarian Riset dan Teknologi Research Permits 40/EXT/SIP/FRP/SM/VII/2015 and 194/SIP/FRP/E5/Dit.KI/VII/2017 and Pusat Penelitian Arkeologi Nasional export permit 2634/H5/TU/2017. For the skeletal material from Liang Bua, Liang Toge and Komodo, the research was undertaken as part of a collaborative project between Universitas Airlangga (Surabaya, Indonesia) and the Max Planck Institute for the Science of Human History (Jena, Germany) under the Kementerian Ristekdikti Research Permit 303/SIP/FRP/E5/Dit.KI/IX/2017 and Pusat Penelitian Arkeologi Nasional export permit 11404H5/TU/2017. Research of the human remains from Jareng Bori were carried out as part of a collaboration between Australian National University and Universitas Gaja Madja under the Kementerian Ristekdikti Research Permit 1209/FRP/E5/Dit.KI/VI/2016.

### *2. Radiocarbon dating.*

Nine new radiocarbon dates for this study (Table S7) were produced at the Curt-Engelhorn-Zentrum Archäometrie gGmbH in Mannheim, Germany. There, collagen from bone and dentin was extracted using a modified Longin method[1] and long molecules removed with ultrafiltration before freeze-drying the product[2]. After the catalytic reduction to graphite the $^{14}C$ content was measured with an AMS-System type MICADAS. The isotopic ratios of $^{14}C/^{12}C$ and $^{13}C/^{12}C$ of samples, standards (Oxalic acid II) and controls were measured simultaneously. The resulting $^{14}C$ dates were normalised to the value $\delta^{13}C=-25‰$[3] and calibrated using the software SwissCal 1.0 (L.Wacker, ETH-Zürich) and the INTCAL13 calibration curve[4]. The Liang Bua individual was dated at the Radiocarbon Dating Laboratory, University of Waikato. It was decalcified in 2%HCl, rinsed and died. Afterwards it was gelatinized for 4 hours at pH 3 with HCL at 90 degrees before ultrafiltration and freeze drying the product. $\delta^{13}C$ values were measured on prepared graphite using the AMS spectrometer and calibrated with OxCal v.4.4 and the INTCAL20 curve[5], accounting for the possible effect of marine $^{14}C$ influence from diet and suggested by the $\delta^{13}C$ and $\delta^{15}N$ values. The Uattamdi individual was dated in the University of Oxford's Radiocarbon Accelerator Unit (ORAU). Bone treatment at the ORAU followed the methods outlined by ref[6]. Bone samples were drilled with tungsten carbide drills and bone powder samples were initially solvent washed due to a suspicion of glue presence on the bone. This comprised a sequence of extraction in acetone, methanol and chloroform. This was then followed by routine ABA pretreatment and ultrafiltration[6]. The ultrafiltered collagen was combusted using an automated carbon and nitrogen elemental analyzer (Carlo Erba EA1108) coupled with a continuous-flow isotope monitoring mass spectrometer (Europa Geo 20/20). $\delta^{15}N$ and $\delta^{13}C$ values, % carbon values, as well as C:N atomic ratios were obtained. We used VPDB as the standard for determining $\delta^{13}C$ values and AIR for the $\delta^{15}N$ equivalent. Graphite was produced from each combusted CO2 sample through a reaction over an iron catalyst in an excess H2 atmosphere at 560°C[7]. AMS radiocarbon measurement was undertaken using the MiCaDaS accelerator and AMS determinations calculated after refs. [3,8]. This sample was corrected for marine reservoir effect using the Mixed_Curves approach in OxCal 4.4 with Marine20 and INTCAL20 curves[5,9] assuming a DR of 0±40 yr in the absence of any local correction data and a contribution of 55.5±10% carbon

derived from a marine protein source. This was determined based on a linear interpolation of the measured $\delta^{13}C$ value (-16.5‰) between assumed endpoints of -21 and -12.5‰ (terrestrial and marine). The Tanjung Pinang individual was dated in the Australian National University. The bone surfaces were cleaned with a dental drill, the sample was crushed and treated with Acetic Acid, rinsed, and dried. Dating was done on bone apatite.
The radiocarbon dates and quality collagen indicators (collagen yields, C/N ratios, %C and %N) are reported in Table S7.

### *3. Archaeological information*

*Aru Manara.* Aru Manara is a limestone cave located about 275 m inland from the southeastern coast of Morotai island, North Maluku Islands, Eastern Indonesia, excavated in 2011. The stratigraphy of the site comprises four deep sediment layers 2 m thick that terminated in the bedrock. The individuals studied here were excavated from Layer 3. The excavations yielded a large number of fragmented human bone (NISP=57,273) and teeth (n=1550), as well as early Metal Age sherds dated to between 2300 and 1800 cal BP, (mainly from Layer 3)[10-12]. The other sherds found at the site (n=504) included 32 red-slipped and 79 decorated specimens. One type of pottery decoration, with linear and curvilinear motifs, are similar to those in contemporary Moluccan assemblages at Tanjung Pinang on Morotai, also analyzed in this study[13]. However, the occurrence of anthropo- and zoomorphic decorations are unique to this site within the Northern Moluccas. The existence of pottery with applied-relief decorations, including a human face and lizard, suggest the development of maritime networks with East Indonesia and Near Oceania from 2700 to 2000 years BP, but is also linked with ceramics from Lapita associated burials in Remote Oceania[14]. The use of large burial jars and box typed jars may indicate possible human interaction between the Northern Moluccas and the Philippines[15]. The skeletal remains from Aru Manara are curated by the Pusat Penelitian Arkeologi Nasional (Jakarta, Indonesia).

*Topogaro 1.* Topogaro 1 is a cave that is part of a larger system of three caves (Topogaro 1-3) and rockshelters (Topogaro 4-7), located about 3.5 km inland from the eastern coast of Central Sulawesi in Morowali District. Topogaro 1 contains more than 30 broken coffins with human skeletal remains and surface finds, including fragments of prehistoric pottery, Chinese and European ceramics, achert flakes (including finely retouched tools), and shell. The presence of a number of flakes and shellfish clearly indicate that the cave had been used as a prehistoric habitation or tool production site, while wooden coffins and the variety of Chinese and European ceramics show the site had been used as a burial site more recently, between 300 and 100 years BP, based on AMS dates acquired from the coffin itself and some associated human teeth. The skeletal remains from Topogaro are curated by the Pusat Penelitian Arkeologi Nasional (Jakarta, Indonesia).

*Gua Uattamdi.* Gua Uattamdi is a limestone rock shelter situated on Kayoa Island in the northern Moluccas (0° 127°20'E). This site has yielded skull fragments associated with jar burial, incised pottery, glass beads and metal fragments, with one individual analysed in this report directly dated to 1697 - 1394 cal BP. The oldest material culture from Uattamdi contained red-slipped Neolithic pottery, dated to c.3300–2500 BP, with connections with the Philippines and eastern Taiwan, as well as other Neolithic sites in Wallacea. This Neolithic layer contained no human remains. The younger incised, impressed and appliqué pottery associated with the analysed skull is paralleled in sites on Morotai, including the two sites of Aru Manara and Tanjung Pinang that are also analysed in this study (ref[13], ch.7). This style of pottery dates to c.2500 BP and later. The skeletal remains from Gua Uattamdi are curated by the Pusat Penelitian Arkeologi Nasional (Jakarta, Indonesia).

*Tanjung Pinang.* Tanjung Pinang is a rock shelter situated on Morotai Island (2°05'N, 128°40'E). There is a direct date for one of the four individuals included in this study (TanjungPinang1) of 2498 - 1687 cal BP. The date was produced from bone apatite in 1993, therefore most likely showing a minimum date (Rachel Wood, ANU Radiocarbon Laboratory, pers. Comm.), and it exhibits a large calibration range. However, the incised decoration of the pottery found at this site is similar to the Metal Age pottery in Gua Uattamdi (ref[13], ch7), supporting a date younger than the Neolithic red-slipped pottery in Uattamdi. Nevertheless, the associated pottery and the apatite analysis allow a possibility that the Tanjung Pinang sample could be as old as 2500 BP. A morphological analysis of the Tanjung Pinang skulls showed Papuan features[16]. The skeletal remains from Tanjung Pinang are curated by the Pusat Penelitian Arkeologi Nasional (Jakarta, Indonesia).

*Liang Bua*. Liang Bua is best known as the cave where *Homo floresiensis* was discovered[17,18]. However, Liang Bua also contains archaeological deposits dating to the Neolithic and Proto-Metallic periods. These later deposits were first excavated in 1965 by T.H. Verhoeven who discovered six skeletons[19,20]. Five skulls from that excavation are curated at Universitas Airlangga and loose petrous bones from Burials 1 and 2 were sampled for the current study. It was later found that both petrous fragments belonged to the crania of Burial 2. The individual included[21] in this study was directly dated to 2588 ± 23 BP (Wk-51763). In both the earlier and later excavations, the burials were found associated with Neolithic and Proto-Metallic material culture and domestic fauna, such as plain and decorated pottery (*periuk* [jars], *kendi*[pitcher/ewer], *buli buli* [little jars], and *tutup* [lid]), flaked adzes, bone tools, pig tusks and a bronze axe[17,20,22]. Bioarchaeological analysis confirmed that 3/3 (100%) of the Liang Bua individuals curated at Universitas Airlangga who had preserved anterior maxillary teeth (including burial 2) had undergone ritual tooth ablation, a practice thought to have been introduced by the Neolithic Austronesian settlers to East Nusa Tenggara[21,23-25].

*Liang Toge.* One individual was analysed in this study from the cave site of Liang Toge on Flores Island, located near Warukia, 1 km south of Lepa, in the Manggarai district of Flores[26]. Excavated by T.H. Verhoeven in the mid 20th century, there is little published information about the human remains from Liang Toge, which are currently curated at Universitas Airlangga, Surabaya. Visual inspection of the Liang Toge skeletal assemblage suggests that these remains may have been (or become after excavation) commingled because there were mixed cranial and post-cranial elements from multiple individuals in the collection and no apparent burial numbers. The petrous bone analyzed for this study was dated to 829 – 728 cal BP (MAMS 40606), in addition to two ribs which provided dates of 1066 – 973 cal BP (MAMS 35084) and 911 – 786 cal BP (MAMS 35085).

*Komodo*. The individual (Komodo I) analysed from Komodo Island, West Manggarai Regency, East Nusa Tenggara (S 08° 32’ 35.9988” E 119° 29’ 21.9876”) was one of two burials found during excavations carried out by Walter Auffenberg (University of Florida, USA) and Putra Sastrawan (Universitas Udayana, Indonesia) in 1969-1970. During their survey for Komodo dragons, Auffenberg and Putra Sastrawan identified 18 archaeological sites across Komodo and the two burials were found at the hill site of Ntodo Leseh. The crania were removed from the primary flexed (or squatting) burials (both with their heads facing west) and they were designated Komodo I and Komodo II[27,28]. Komodo I was buried 16 cm below the surface and had a circle of flat stones marking the grave. Komodo II was buried 19 cm below the surface and had a triangle of flat stones marking the grave. From comparisons of burial type with other prehistoric interments on Flores and East Java, in addition to material culture (stone flakes and blades and Neolithic hand axes) found at the other sites on Komodo, ref. [21] proposed that the Komodo burials were between 3000-5000 years old. However, recent direct dating of the petrous bone from Komodo I provided a much more modern date of 687 - 661 cal BP (MAMS 40602) indicating that this individual lived during the period when East Nusa Tenggara was under the rule of the Majapahit Empire[29]. The skeletal remains from Jareng Bori are curated at Universitas Airlangga (Surabaya, Indonesia).

*Jareng Bori.* The site of Jareng Bori, Pantar island, East Nusa Tenggara (S 08° 15’51.7 E 124°17’55.4) was excavated in 2016 as part of a joint research project between researchers from the Australian National University (Australia) and Universitas Gaja Madja (Indonesia). The small rock shelter (40 m2 living floor) was located beside a large boulder at the base of a cliff on a coastal beach flat approximately 120 m from the current shoreline. A 1 x 1 m test pit (test pit A) was excavated and seven stratigraphic layers were identified at the site. The period of site use was dated between the Metal Age, *ca* 1800 BP, and the late Historic Period. A site report detailing the full excavation and results of the dating, faunal and material culture analyses is available[30]. At Jareng Bori, a poorly preserved human burial (analyzed for the current study) was found cut into the upper layers with a skeleton in a flexed position in the south area of the test pit. Only the upper half (thorax, upper limbs and skull) of the skeleton was excavated and the remainder was left *in situ* because it extended into the southern baulk of the test pit. The skeletal remains were analysed at Australian

National University and a full bioarchaeology report is forthcoming. Of the 21 charcoal dates analysed from the site, four were associated with the burial (ANU 53127, ANU 53130, ANU 53129, and ANU136) and ranged between 0-429 cal BP[30]. The direct date of the petrous bone for this burial provided results of 527 – 498 cal BP (MAMS 40607), indicating that this individual lived during the rule of the Majahapit Empire in the region, directly before the first Portuguese contact with the area at the beginning of the 16th century AD[29]. Tooth modifications in the form of labial filing of the maxillary incisors was found on the dentition of the Jareng Bori individual. This type of tooth modification has been observed on skeletons dating to a similar time period in Java, Bali, Sumba and Flores. Tooth filing has been suggested as a cultural tradition that is a unique characteristic of later inhabitants of the region[23,35,31], whereas tooth ablation (in the form of the removal of the maxillary lateral incisors and canines) has been interpreted as an earlier Neolithic/Austronesian ritual practiced across East Nusa Tenggara *ca.* 3000 - 2000 BP[23-25]. The skeletal remains from Jareng Bori are curated at Universitas Airlangga (Surabaya, Indonesia).

**Figure S1 - PCA of publicly available whole genome data merged with Human Origins genotype data and Affymetrix Axiom Genome-Wide Human genotype data.** Ancient individuals (shown with a black contour) are projected and their fill color matches the color of present-day samples from the same geographic area. The location of the ancient individuals newly presented in this study is shown in the map on the right panel.

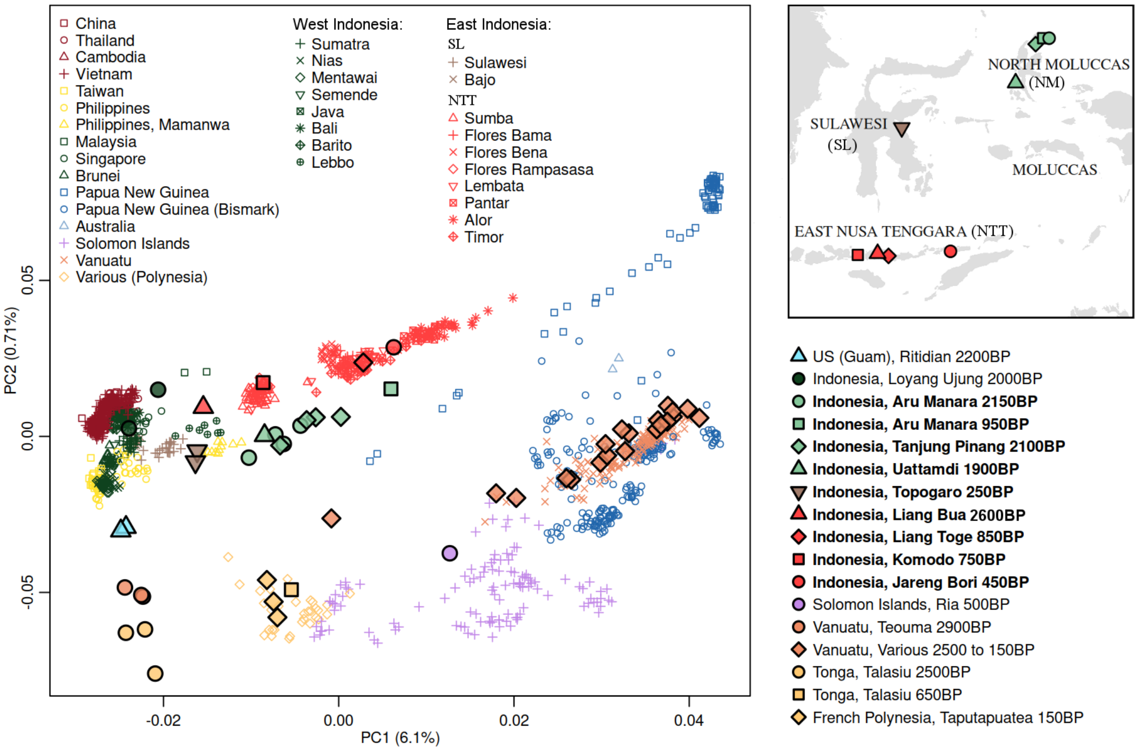

**Figure S2 - DyStruct results**. A-B Conditional log likelihood (CLL) values across K = 2 to K = 15 obtained for 25 independent runs (grey circles) using datasets that merge publicly available whole genome data and Human Origins genotype data, with Affymetrix 6.0 (A) and Affymetrix Axiom Genome-Wide Human (B) genotype data. Black dots show the average CLL per K. The yellow bar indicates the value of K at which the CLL starts to plateau. C-D ADMIXTURE-like barplots for the highlighted K value of each respective dataset: K= 13 for the Affymetrix 6.0 merge (C); and K = 9 for the Affymetrix Axiom Genome-Wide Human merge (D).

A)

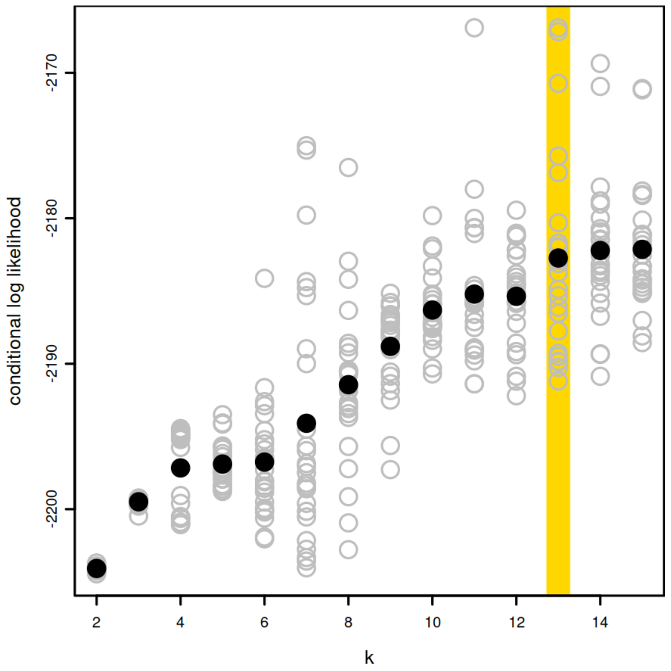


B)

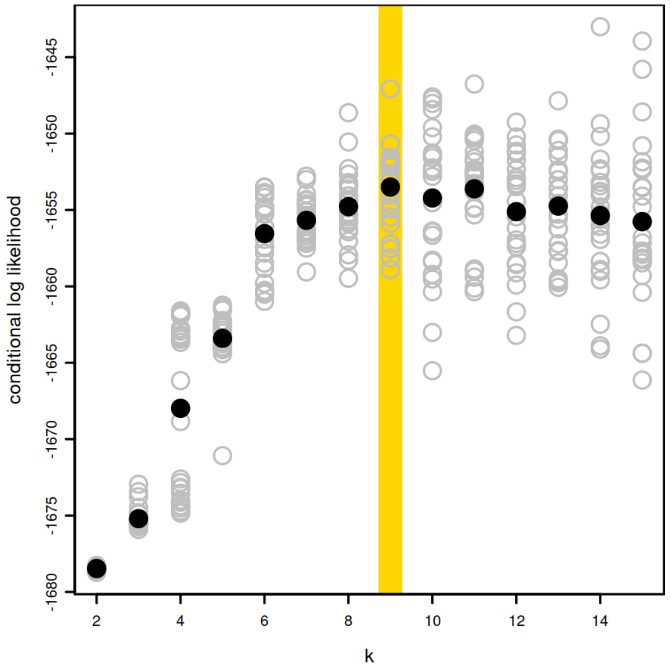

C)

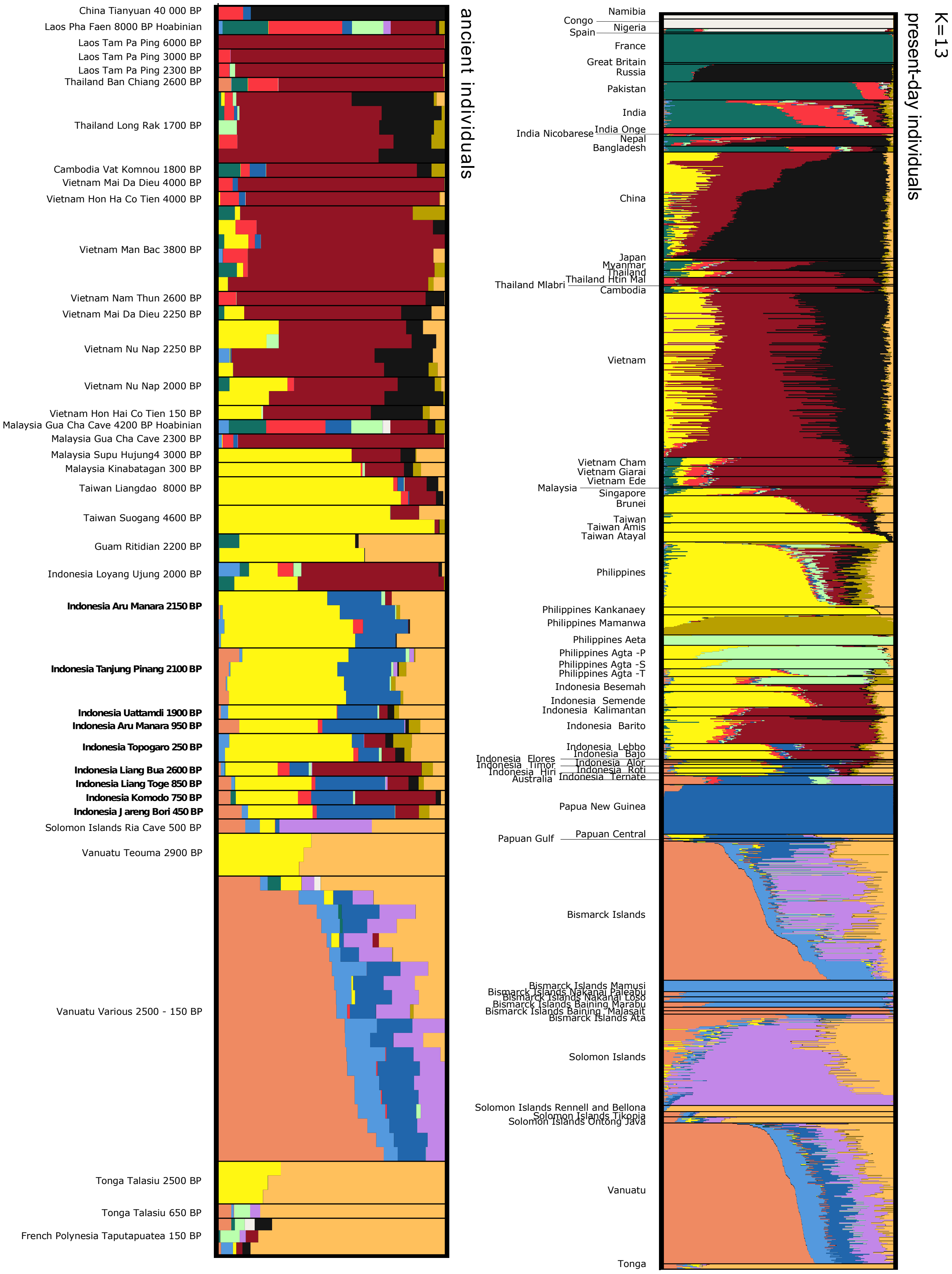
China Tianyuan 40 000 BP
Laos Pha Faen 8000 BP Hoabinian
Laos Tam Pa Ping 6000 BP
Laos Tam Pa Ping 3000 BP
Laos Tam Pa Ping 2300 BP
Thailand Ban Chiang 2600 BP
Thailand Long Rak 1700 BP
Cambodia Vat Komnou 1800 BP
Vietnam Mai Da Dieu 4000 BP
Vietnam Hon Ha Co Tien 4000 BP
Vietnam Man Bac 3800 BP
Vietnam Nam Thun 2600 BP
Vietnam Mai Da Dieu 2250 BP
Vietnam Nu Nap 2250 BP
Vietnam Nu Nap 2000 BP
Vietnam Hon Hai Co Tien 150 BP
Malaysia Gua Cha Cave 4200 BP Hoabinian
Malaysia Gua Cha Cave 2300 BP
Malaysia Supu Hujung4 3000 BP
Malaysia Kinabatagan 300 BP
Taiwan Liangdao 8000 BP
Taiwan Suogang 4600 BP
Guam Ritidian 2200 BP
Indonesia Loyang Ujung 2000 BP
Indonesia Aru Manara 2150 BP
Indonesia Tanjung Pinang 2100 BP
Indonesia Uattamdi 1900 BP
Indonesia Aru Manara 950 BP
Indonesia Topogaro 250 BP
Indonesia Liang Bua 2600 BP
Indonesia Liang Toge 850 BP
Indonesia Komodo 750 BP
Indonesia Jareng Bori 450 BP
Solomon Islands Ria Cave 500 BP
Vanuatu Teouma 2900 BP
Vanuatu Various 2500 - 150 BP
Tonga Talasiu 2500 BP
Tonga Talasiu 650 BP
French Polynesia Taputapuatea 150 BP
ancient individuals
Namibia
Congo
Nigeria
Spain
France
Great Britain
Russia
Pakistan
India
India Onge
India Nicobarese
Nepal
Bangladesh
China
Japan
Myanmar
Thailand
Thailand Htin Mal
Thailand Mlabri
Cambodia
Vietnam
Vietnam Cham
Vietnam Giarai
Vietnam Ede
Malaysia
Singapore
Brunei
Taiwan
Taiwan Amis
Taiwan Atayal
Philippines
Philippines Kankanaey
Philippines Mamanwa
Philippines Aeta
Philippines Agta -P
Philippines Agta -S
Philippines Agta -T
Indonesia Besemah
Indonesia Semende
Indonesia Kalimantan
Indonesia Barito
Indonesia Lebbo
Indonesia Bajo
Indonesia Flores
Indonesia Timor
Indonesia Alor
Indonesia Hiri
Indonesia Roti
Australia
Indonesia Ternate
Papua New Guinea
Papuan Gulf
Papuan Central
Bismarck Islands
Bismarck Islands Mamusi
Bismarck Islands Nakanai Paleabu
Bismarck Islands Nakanai Loso
Bismarck Islands Baining Marabu
Bismarck Islands Baining Malasait
Bismarck Islands Ata
Solomon Islands
Solomon Islands Rennell and Bellona
Solomon Islands Tikopia
Solomon Islands Ontong Java
Vanuatu
Tonga
present-day individuals
K=13

D)

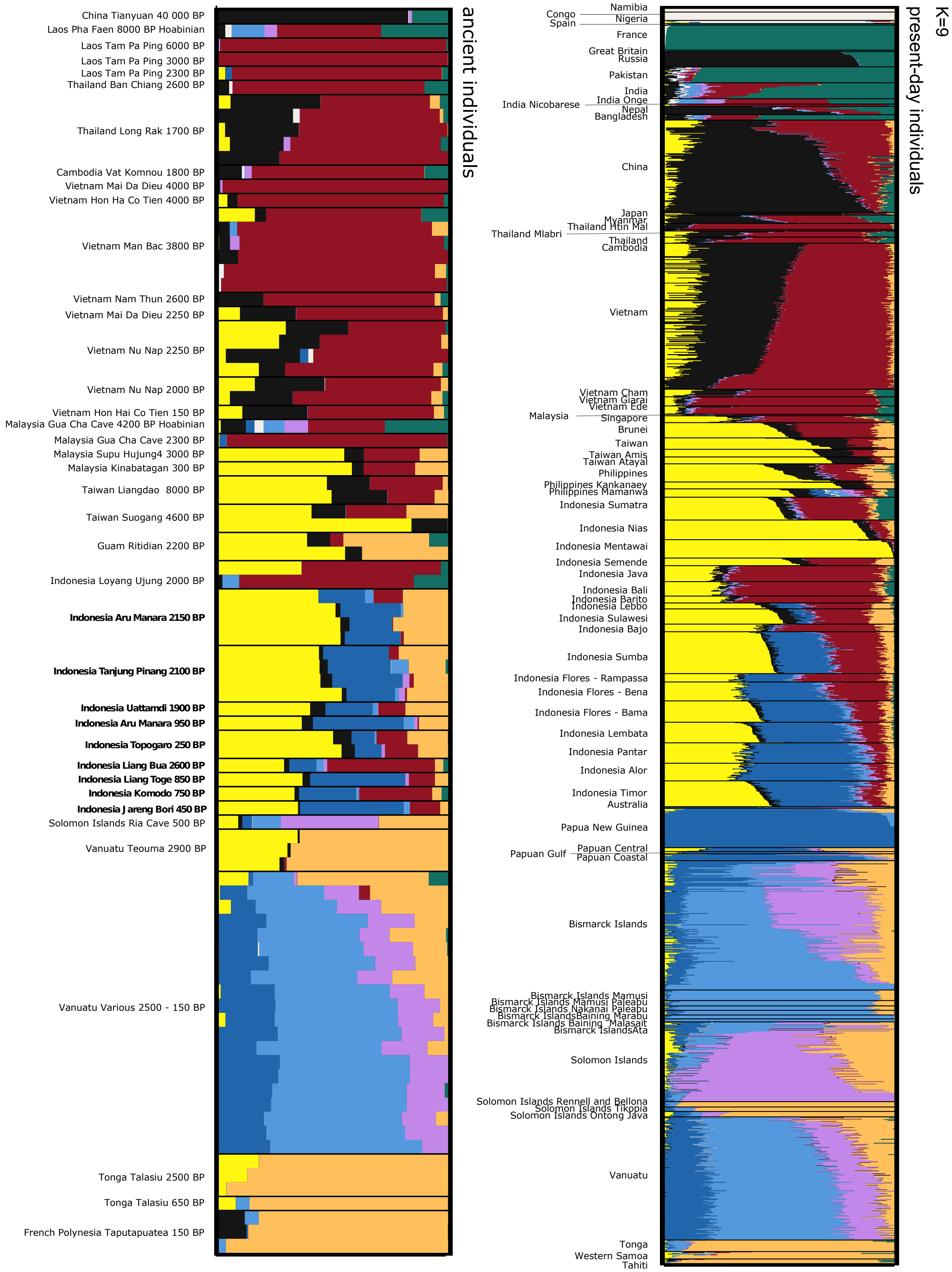

ancient individuals
China Tianyuan 40 000 BP
Laos Pha Faen 8000 BP Hoabinian
Laos Tam Pa Ping 6000 BP
Laos Tam Pa Ping 3000 BP
Laos Tam Pa Ping 2300 BP
Thailand Ban Chiang 2600 BP
Thailand Long Rak 1700 BP
Cambodia Vat Komnou 1800 BP
Vietnam Mai Da Dieu 4000 BP
Vietnam Hon Ha Co Tien 4000 BP
Vietnam Man Bac 3800 BP
Vietnam Nam Thun 2600 BP
Vietnam Mai Da Dieu 2250 BP
Vietnam Nu Nap 2250 BP
Vietnam Nu Nap 2000 BP
Vietnam Hon Hai Co Tien 150 BP
Malaysia Gua Cha Cave 4200 BP Hoabinian
Malaysia Gua Cha Cave 2300 BP
Malaysia Supu Hujung4 3000 BP
Malaysia Kinabatagan 300 BP
Taiwan Liangdao 8000 BP
Taiwan Suogang 4600 BP
Guam Ritidian 2200 BP
Indonesia Loyang Ujung 2000 BP
Indonesia Aru Manara 2150 BP
Indonesia Tanjung Pinang 2100 BP
Indonesia Uattamdi 1900 BP
Indonesia Aru Manara 950 BP
Indonesia Topogaro 250 BP
Indonesia Liang Bua 2600 BP
Indonesia Liang Toge 850 BP
Indonesia Komodo 750 BP
Indonesia Jareng Bori 450 BP
Solomon Islands Ria Cave 500 BP
Vanuatu Teouma 2900 BP
Vanuatu Various 2500 - 150 BP
Tonga Talasiu 2500 BP
Tonga Talasiu 650 BP
French Polynesia Taputapuatea 150 BP
K=9
present-day individuals
Namibia
Congo
Nigeria
Spain
France
Great Britain
Russia
Pakistan
India
India Onge
India Nicobarese
Nepal
Bangladesh
China
Japan
Myanmar
Thailand Htin Mal
Thailand Mlabri
Thailand
Cambodia
Vietnam
Vietnam Cham
Vietnam Giarai
Vietnam Ede
Malaysia
Singapore
Brunei
Taiwan
Taiwan Amis
Taiwan Atayal
Philippines
Philippines Kankanaey
Philippines Mamanwa
Indonesia Sumatra
Indonesia Nias
Indonesia Mentawai
Indonesia Semende
Indonesia Java
Indonesia Bali
Indonesia Barito
Indonesia Lebbo
Indonesia Sulawesi
Indonesia Bajo
Indonesia Sumba
Indonesia Flores - Rampassa
Indonesia Flores - Bena
Indonesia Flores - Bama
Indonesia Lembata
Indonesia Pantar
Indonesia Alor
Indonesia Timor
Australia
Papua New Guinea
Papuan Gulf
Papuan Central
Papuan Coastal
Bismarck Islands
Bismarck Islands Mamusi
Bismarck Islands Mamusi Paleabu
Bismarck Islands Nakanai Paleabu
Bismarck IslandsBaining Marabu
Bismarck Islands Baining Malasait
Bismarck IslandsAta
Solomon Islands
Solomon Islands Rennell and Bellona
Solomon Islands Tikopia
Solomon Islands Ontong Java
Vanuatu
Tonga
Western Samoa
Tahiti

**Figure S3 - Representation of two Austronesian-related components**. The frequencies of the "yellow" and "mango" components identified in Supplementary Figure 1C were normalized to sum to 1.

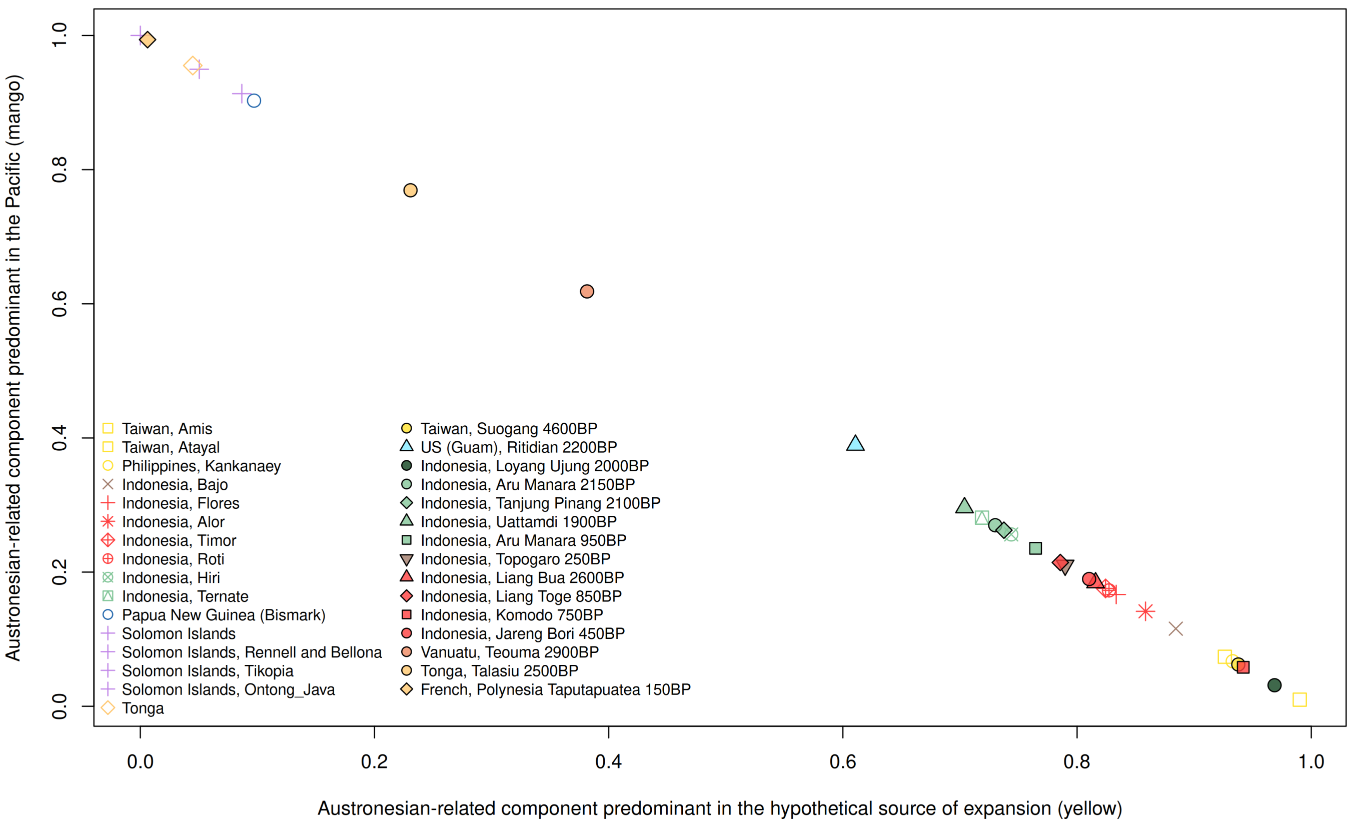

**Figure S4 - $f_4$-statistic of the form $f_4$ *(Mbuti, ancient Wallacean; Amis, test)* computed for each ancient group from Wallacea separately.** The test groups are shown in the y-axis and include ancient and present-day groups from mainland Asia, Island southeast Asia, and the Pacific that have no discernible Papuan-related ancestry. Bars indicate two standard errors in both directions. Values in green are not significantly different from zero (|Z| < 2), whereas values in red are significantly different from zero (|Z| > 2).

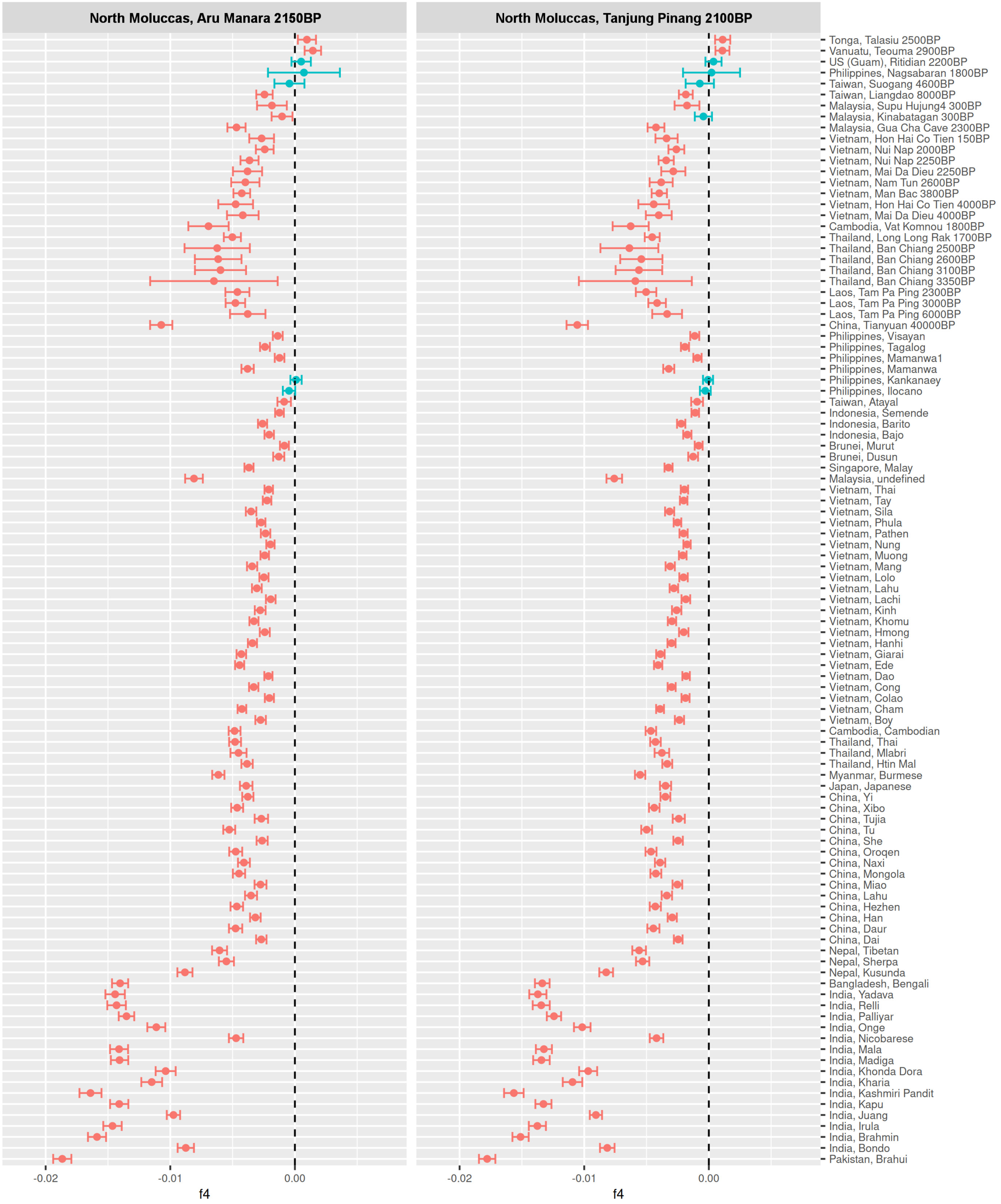

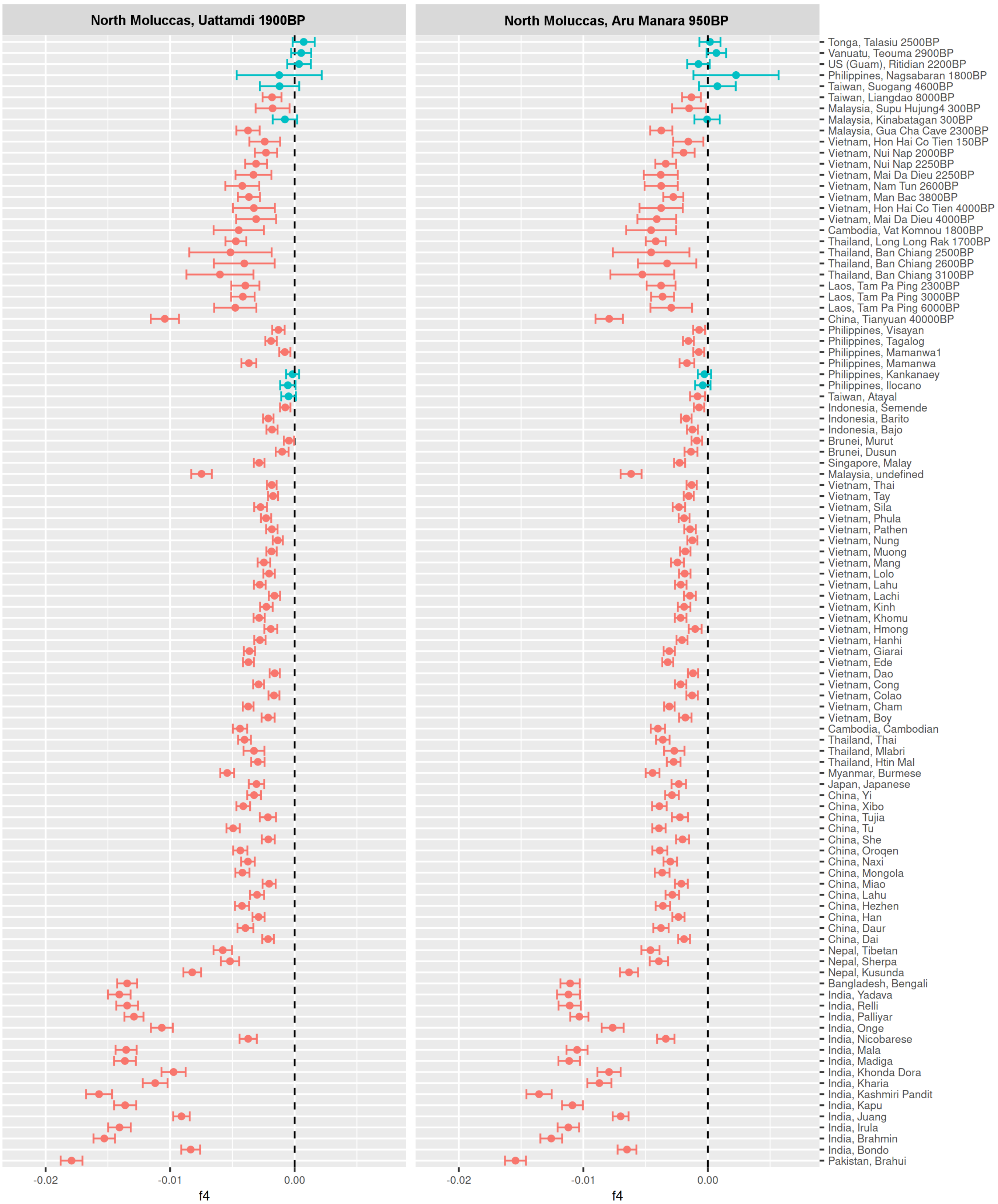

North Moluccas, Uattamdi 1900BP
North Moluccas, Aru Manara 950BP
Tonga, Talasiu 2500BP
Vanuatu, Teouma 2900BP
US (Guam), Ritidian 2200BP
Philippines, Nagsabaran 1800BP
Taiwan, Suogang 4600BP
Taiwan, Liangdao 8000BP
Malaysia, Supu Hujung4 300BP
Malaysia, Kinabatagan 300BP
Malaysia, Gua Cha Cave 2300BP
Vietnam, Hon Hai Co Tien 150BP
Vietnam, Nui Nap 2000BP
Vietnam, Nui Nap 2250BP
Vietnam, Mai Da Dieu 2250BP
Vietnam, Nam Tun 2600BP
Vietnam, Man Bac 3800BP
Vietnam, Hon Hai Co Tien 4000BP
Vietnam, Mai Da Dieu 4000BP
Cambodia, Vat Komnou 1800BP
Thailand, Long Long Rak 1700BP
Thailand, Ban Chiang 2500BP
Thailand, Ban Chiang 2600BP
Thailand, Ban Chiang 3100BP
Laos, Tam Pa Ping 2300BP
Laos, Tam Pa Ping 3000BP
Laos, Tam Pa Ping 6000BP
China, Tianyuan 40000BP
Philippines, Visayan
Philippines, Tagalog
Philippines, Mamanwa1
Philippines, Mamanwa
Philippines, Kankanaey
Philippines, Ilocano
Taiwan, Atayal
Indonesia, Semende
Indonesia, Barito
Indonesia, Bajo
Brunei, Murut
Brunei, Dusun
Singapore, Malay
Malaysia, undefined
Vietnam, Thai
Vietnam, Tay
Vietnam, Sila
Vietnam, Phula
Vietnam, Pathen
Vietnam, Nung
Vietnam, Muong
Vietnam, Mang
Vietnam, Lolo
Vietnam, Lahu
Vietnam, Lachi
Vietnam, Kinh
Vietnam, Khomu
Vietnam, Hmong
Vietnam, Hanhi
Vietnam, Giarai
Vietnam, Ede
Vietnam, Dao
Vietnam, Cong
Vietnam, Colao
Vietnam, Cham
Vietnam, Boy
Cambodia, Cambodian
Thailand, Thai
Thailand, Mlabri
Thailand, Htin Mal
Myanmar, Burmese
Japan, Japanese
China, Yi
China, Xibo
China, Tujia
China, Tu
China, She
China, Oroqen
China, Naxi
China, Mongola
China, Miao
China, Lahu
China, Hezhen
China, Han
China, Daur
China, Dai
Nepal, Tibetan
Nepal, Sherpa
Nepal, Kusunda
Bangladesh, Bengali
India, Yadava
India, Relli
India, Palliyar
India, Onge
India, Nicobarese
India, Mala
India, Madiga
India, Khonda Dora
India, Kharia
India, Kashmiri Pandit
India, Kapu
India, Juang
India, Irula
India, Brahmin
India, Bondo
Pakistan, Brahui
-0.02
-0.01
0.00
f4

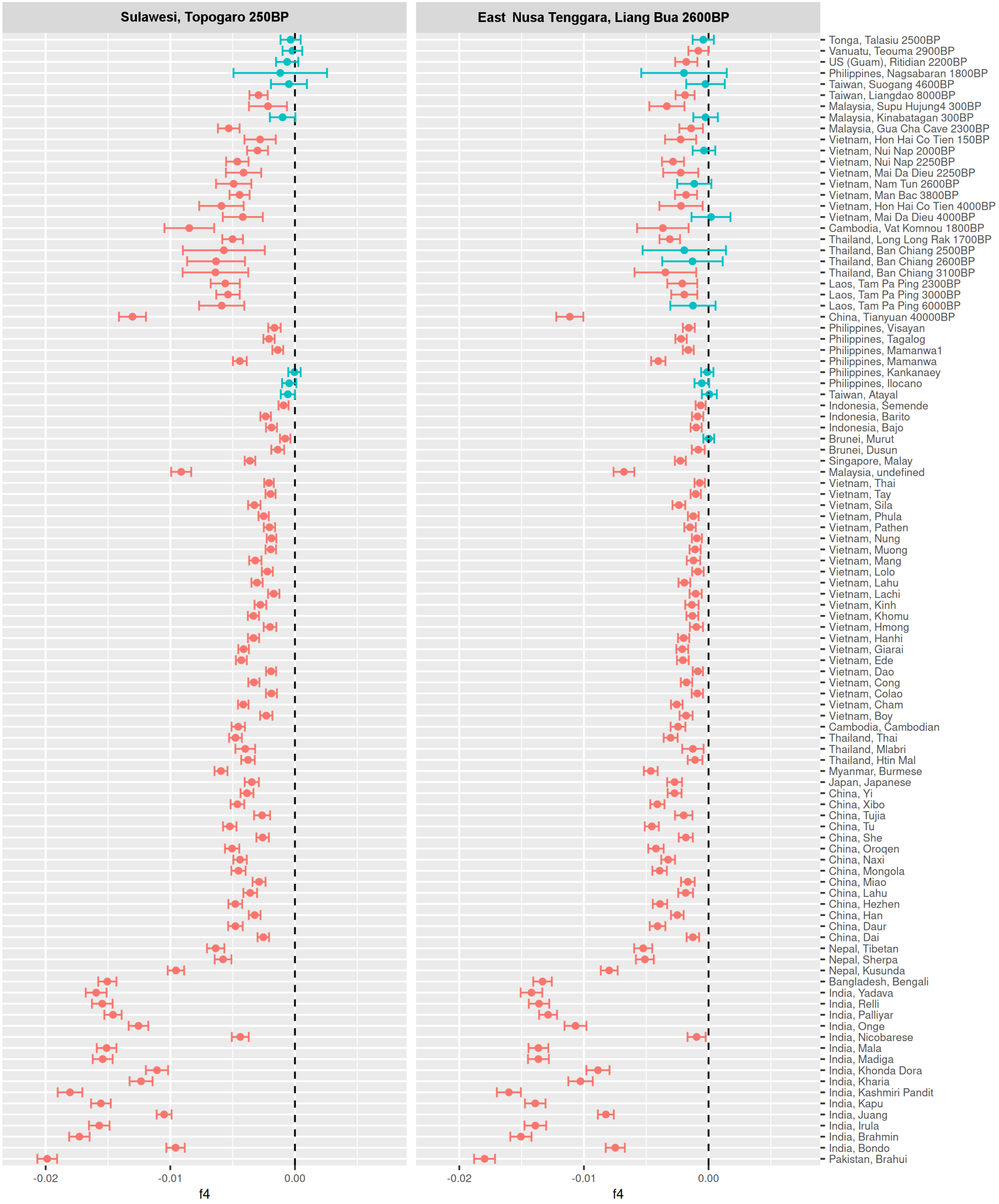

Sulawesi, Topogaro 250BP
East Nusa Tenggara, Liang Bua 2600BP
Tonga, Talasiu 2500BP
Vanuatu, Teouma 2900BP
US (Guam), Ritidian 2200BP
Philippines, Nagsabaran 1800BP
Taiwan, Suogang 4600BP
Taiwan, Liangdao 8000BP
Malaysia, Supu Hujung4 300BP
Malaysia, Kinabatagan 300BP
Malaysia, Gua Cha Cave 2300BP
Vietnam, Hon Hai Co Tien 150BP
Vietnam, Nui Nap 2000BP
Vietnam, Nui Nap 2250BP
Vietnam, Mai Da Dieu 2250BP
Vietnam, Nam Tun 2600BP
Vietnam, Man Bac 3800BP
Vietnam, Hon Hai Co Tien 4000BP
Vietnam, Mai Da Dieu 4000BP
Cambodia, Vat Komnou 1800BP
Thailand, Long Long Rak 1700BP
Thailand, Ban Chiang 2500BP
Thailand, Ban Chiang 2600BP
Thailand, Ban Chiang 3100BP
Laos, Tam Pa Ping 2300BP
Laos, Tam Pa Ping 3000BP
Laos, Tam Pa Ping 6000BP
China, Tianyuan 40000BP
Philippines, Visayan
Philippines, Tagalog
Philippines, Mamanwa1
Philippines, Mamanwa
Philippines, Kankanaey
Philippines, Ilocano
Taiwan, Atayal
Indonesia, Semende
Indonesia, Barito
Indonesia, Bajo
Brunei, Murut
Brunei, Dusun
Singapore, Malay
Malaysia, undefined
Vietnam, Thai
Vietnam, Tay
Vietnam, Sila
Vietnam, Phula
Vietnam, Pathen
Vietnam, Nung
Vietnam, Muong
Vietnam, Mang
Vietnam, Lolo
Vietnam, Lahu
Vietnam, Lachi
Vietnam, Kinh
Vietnam, Khomu
Vietnam, Hmong
Vietnam, Hanhi
Vietnam, Giarai
Vietnam, Ede
Vietnam, Dao
Vietnam, Cong
Vietnam, Colao
Vietnam, Cham
Vietnam, Boy
Cambodia, Cambodian
Thailand, Thai
Thailand, Mlabri
Thailand, Htin Mal
Myanmar, Burmese
Japan, Japanese
China, Yi
China, Xibo
China, Tujia
China, Tu
China, She
China, Oroqen
China, Naxi
China, Mongola
China, Miao
China, Lahu
China, Hezhen
China, Han
China, Daur
China, Dai
Nepal, Tibetan
Nepal, Sherpa
Nepal, Kusunda
Bangladesh, Bengali
India, Yadava
India, Relli
India, Palliyar
India, Onge
India, Nicobarese
India, Mala
India, Madiga
India, Khonda Dora
India, Kharia
India, Kashmiri Pandit
India, Kapu
India, Juang
India, Irula
India, Brahmin
India, Bondo
Pakistan, Brahui
-0.02
-0.01
0.00
f4

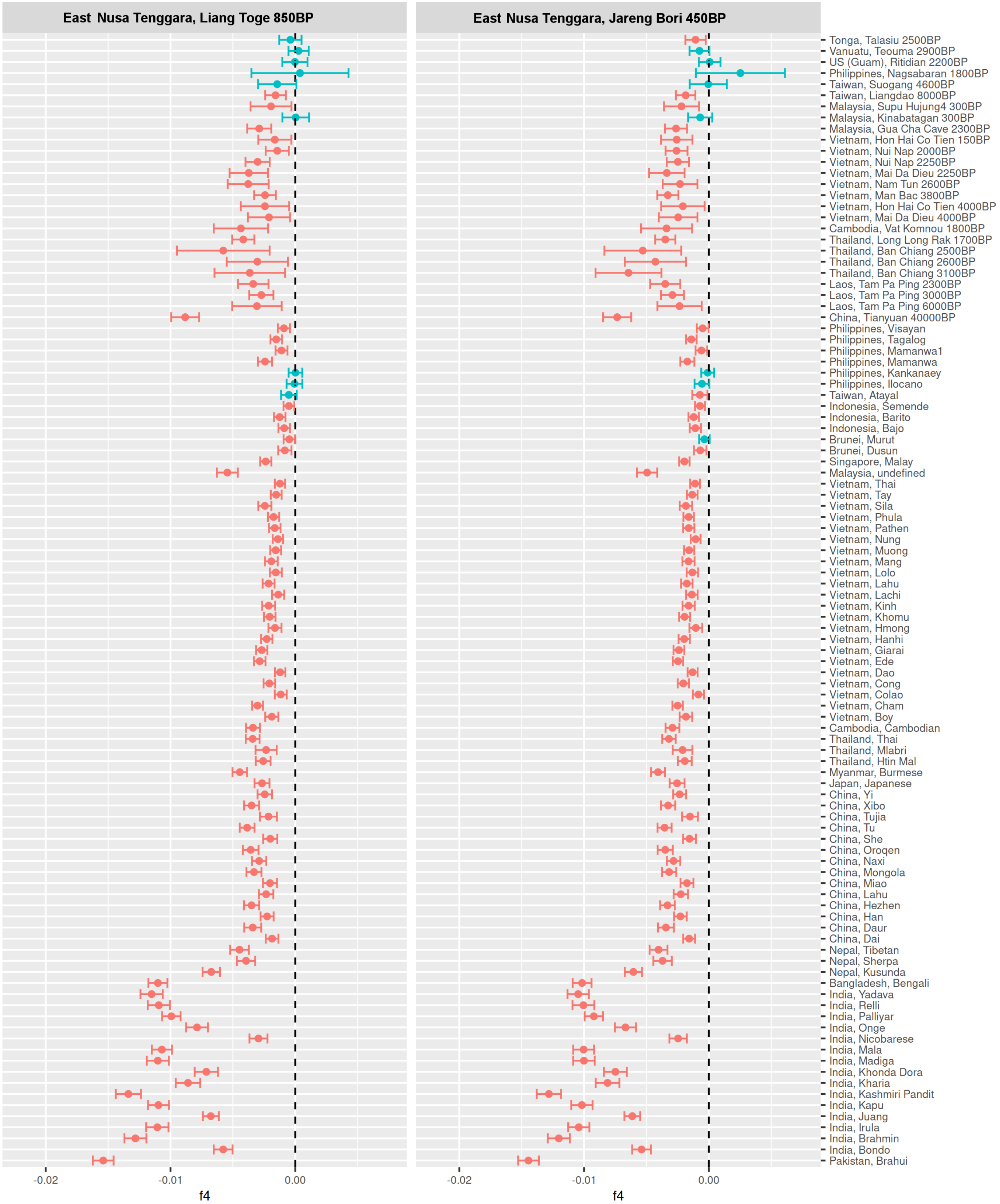

East Nusa Tenggara, Liang Toge 850BP
East Nusa Tenggara, Jareng Bori 450BP
Tonga, Talasiu 2500BP
Vanuatu, Teouma 2900BP
US (Guam), Ritidian 2200BP
Philippines, Nagsabaran 1800BP
Taiwan, Suogang 4600BP
Taiwan, Liangdao 8000BP
Malaysia, Supu Hujung4 300BP
Malaysia, Kinabatagan 300BP
Malaysia, Gua Cha Cave 2300BP
Vietnam, Hon Hai Co Tien 150BP
Vietnam, Nui Nap 2000BP
Vietnam, Nui Nap 2250BP
Vietnam, Mai Da Dieu 2250BP
Vietnam, Nam Tun 2600BP
Vietnam, Man Bac 3800BP
Vietnam, Hon Hai Co Tien 4000BP
Vietnam, Mai Da Dieu 4000BP
Cambodia, Vat Komnou 1800BP
Thailand, Long Long Rak 1700BP
Thailand, Ban Chiang 2500BP
Thailand, Ban Chiang 2600BP
Thailand, Ban Chiang 3100BP
Laos, Tam Pa Ping 2300BP
Laos, Tam Pa Ping 3000BP
Laos, Tam Pa Ping 6000BP
China, Tianyuan 40000BP
Philippines, Visayan
Philippines, Tagalog
Philippines, Mamanwa1
Philippines, Mamanwa
Philippines, Kankanaey
Philippines, Ilocano
Taiwan, Atayal
Indonesia, Semende
Indonesia, Barito
Indonesia, Bajo
Brunei, Murut
Brunei, Dusun
Singapore, Malay
Malaysia, undefined
Vietnam, Thai
Vietnam, Tay
Vietnam, Sila
Vietnam, Phula
Vietnam, Pathen
Vietnam, Nung
Vietnam, Muong
Vietnam, Mang
Vietnam, Lolo
Vietnam, Lahu
Vietnam, Lachi
Vietnam, Kinh
Vietnam, Khomu
Vietnam, Hmong
Vietnam, Hanhi
Vietnam, Giarai
Vietnam, Ede
Vietnam, Dao
Vietnam, Cong
Vietnam, Colao
Vietnam, Cham
Vietnam, Boy
Cambodia, Cambodian
Thailand, Thai
Thailand, Mlabri
Thailand, Htin Mal
Myanmar, Burmese
Japan, Japanese
China, Yi
China, Xibo
China, Tujia
China, Tu
China, She
China, Oroqen
China, Naxi
China, Mongola
China, Miao
China, Lahu
China, Hezhen
China, Han
China, Daur
China, Dai
Nepal, Tibetan
Nepal, Sherpa
Nepal, Kusunda
Bangladesh, Bengali
India, Yadava
India, Relli
India, Palliyar
India, Onge
India, Nicobarese
India, Mala
India, Madiga
India, Khonda Dora
India, Kharia
India, Kashmiri Pandit
India, Kapu
India, Juang
India, Irula
India, Brahmin
India, Bondo
Pakistan, Brahui
-0.02
-0.01
0.00
f4
-0.02
-0.01
0.00
f4

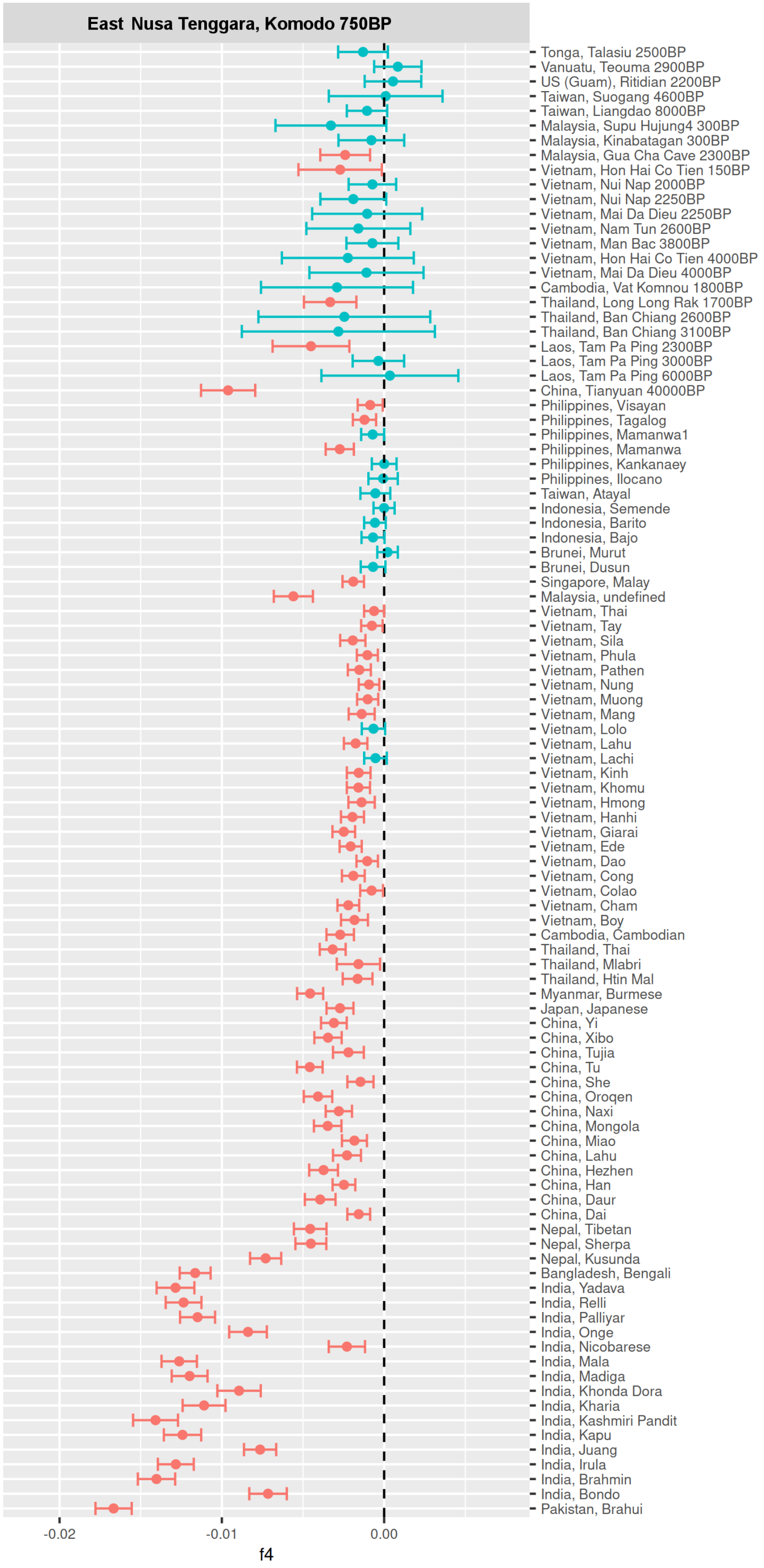
East Nusa Tenggara, Komodo 750BP
Tonga, Talasiu 2500BP
Vanuatu, Teouma 2900BP
US (Guam), Ritidian 2200BP
Taiwan, Suogang 4600BP
Taiwan, Liangdao 8000BP
Malaysia, Supu Hujung4 300BP
Malaysia, Kinabatagan 300BP
Malaysia, Gua Cha Cave 2300BP
Vietnam, Hon Hai Co Tien 150BP
Vietnam, Nui Nap 2000BP
Vietnam, Nui Nap 2250BP
Vietnam, Mai Da Dieu 2250BP
Vietnam, Nam Tun 2600BP
Vietnam, Man Bac 3800BP
Vietnam, Hon Hai Co Tien 4000BP
Vietnam, Mai Da Dieu 4000BP
Cambodia, Vat Komnou 1800BP
Thailand, Long Long Rak 1700BP
Thailand, Ban Chiang 2600BP
Thailand, Ban Chiang 3100BP
Laos, Tam Pa Ping 2300BP
Laos, Tam Pa Ping 3000BP
Laos, Tam Pa Ping 6000BP
China, Tianyuan 40000BP
Philippines, Visayan
Philippines, Tagalog
Philippines, Mamanwa1
Philippines, Mamanwa
Philippines, Kankanaey
Philippines, Ilocano
Taiwan, Atayal
Indonesia, Semende
Indonesia, Barito
Indonesia, Bajo
Brunei, Murut
Brunei, Dusun
Singapore, Malay
Malaysia, undefined
Vietnam, Thai
Vietnam, Tay
Vietnam, Sila
Vietnam, Phula
Vietnam, Pathen
Vietnam, Nung
Vietnam, Muong
Vietnam, Mang
Vietnam, Lolo
Vietnam, Lahu
Vietnam, Lachi
Vietnam, Kinh
Vietnam, Khomu
Vietnam, Hmong
Vietnam, Hanhi
Vietnam, Giarai
Vietnam, Ede
Vietnam, Dao
Vietnam, Cong
Vietnam, Colao
Vietnam, Cham
Vietnam, Boy
Cambodia, Cambodian
Thailand, Thai
Thailand, Mlabri
Thailand, Htin Mal
Myanmar, Burmese
Japan, Japanese
China, Yi
China, Xibo
China, Tujia
China, Tu
China, She
China, Oroqen
China, Naxi
China, Mongola
China, Miao
China, Lahu
China, Hezhen
China, Han
China, Daur
China, Dai
Nepal, Tibetan
Nepal, Sherpa
Nepal, Kusunda
Bangladesh, Bengali
India, Yadava
India, Relli
India, Palliyar
India, Onge
India, Nicobarese
India, Mala
India, Madiga
India, Khonda Dora
India, Kharia
India, Kashmiri Pandit
India, Kapu
India, Juang
India, Irula
India, Brahmin
India, Bondo
Pakistan, Brahui
-0.02
-0.01
0.00
f4

**Figure S5 - Biplots showing the results of pairs of $f_4$-statistics of the form: $f_4$(Mbuti, test; New Guinea Highlanders, ancient Wallacea).** The test groups, shown on the x-y axis label, include present-day groups from mainland Asia, Island southeast Asia, and Oceania that have no discernible Papuan-related ancestry based on the DyStruct analysis. Grey bars show two standard errors in each direction. Linear regression lines for the North Moluccas and East Nusa Tenggara individuals are shown in green and red, respectively.

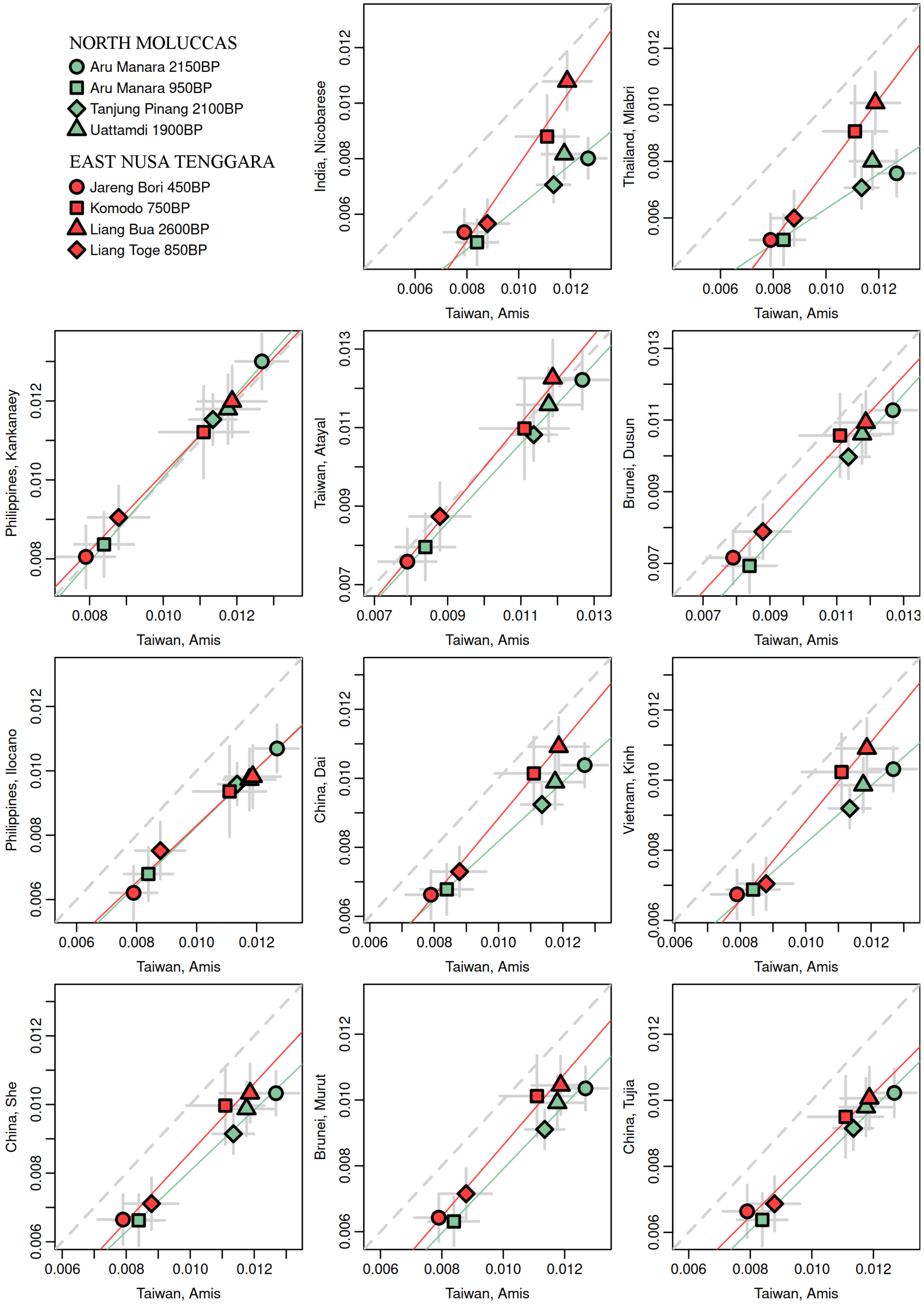

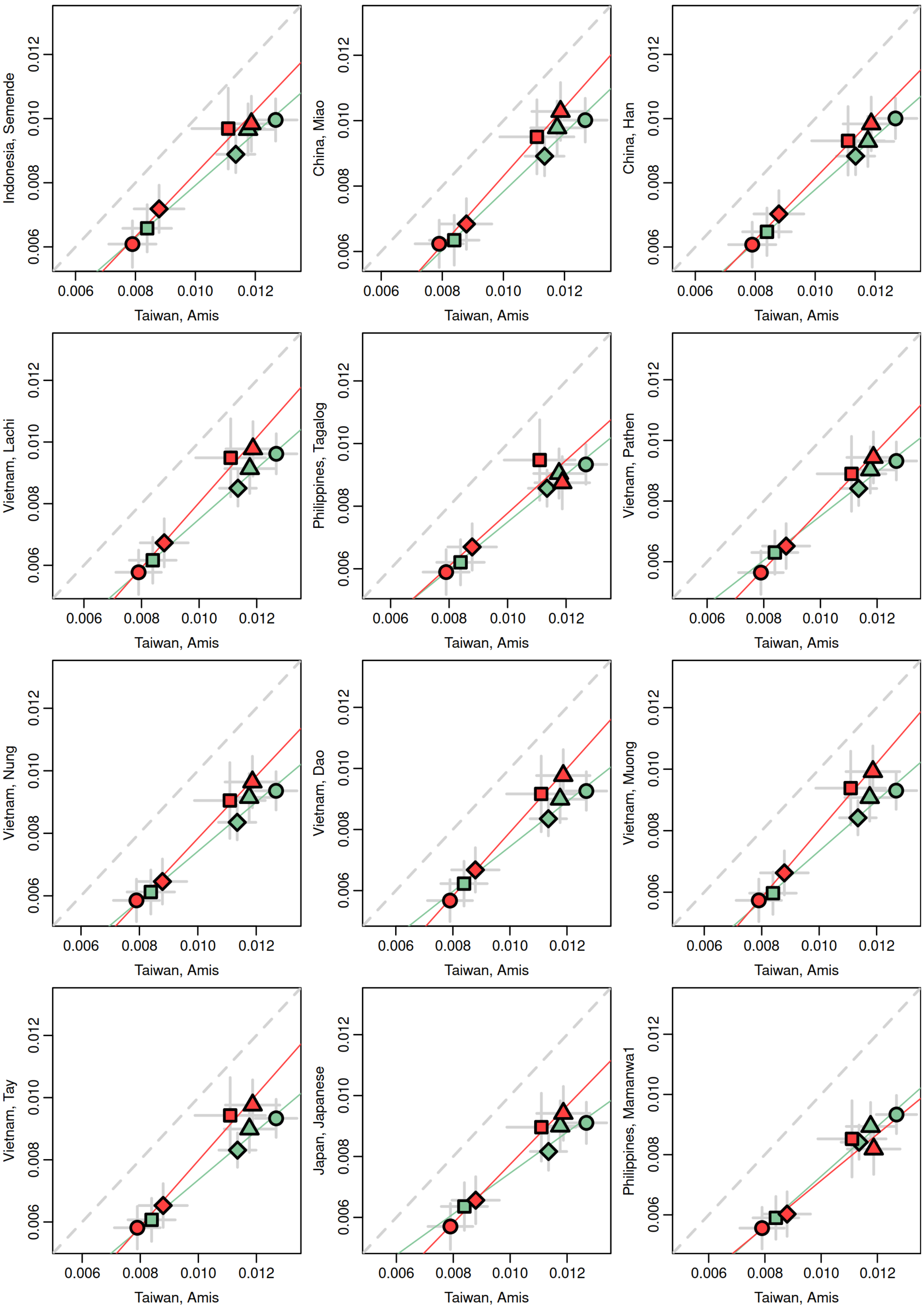
Indonesia, Semende
China, Miao
China, Han
Vietnam, Lachi
Philippines, Tagalog
Vietnam, Pathen
Vietnam, Nung
Vietnam, Dao
Vietnam, Muong
Vietnam, Tay
Japan, Japanese
Philippines, Mamanwa1
0.006
0.008
0.010
0.012
Taiwan, Amis

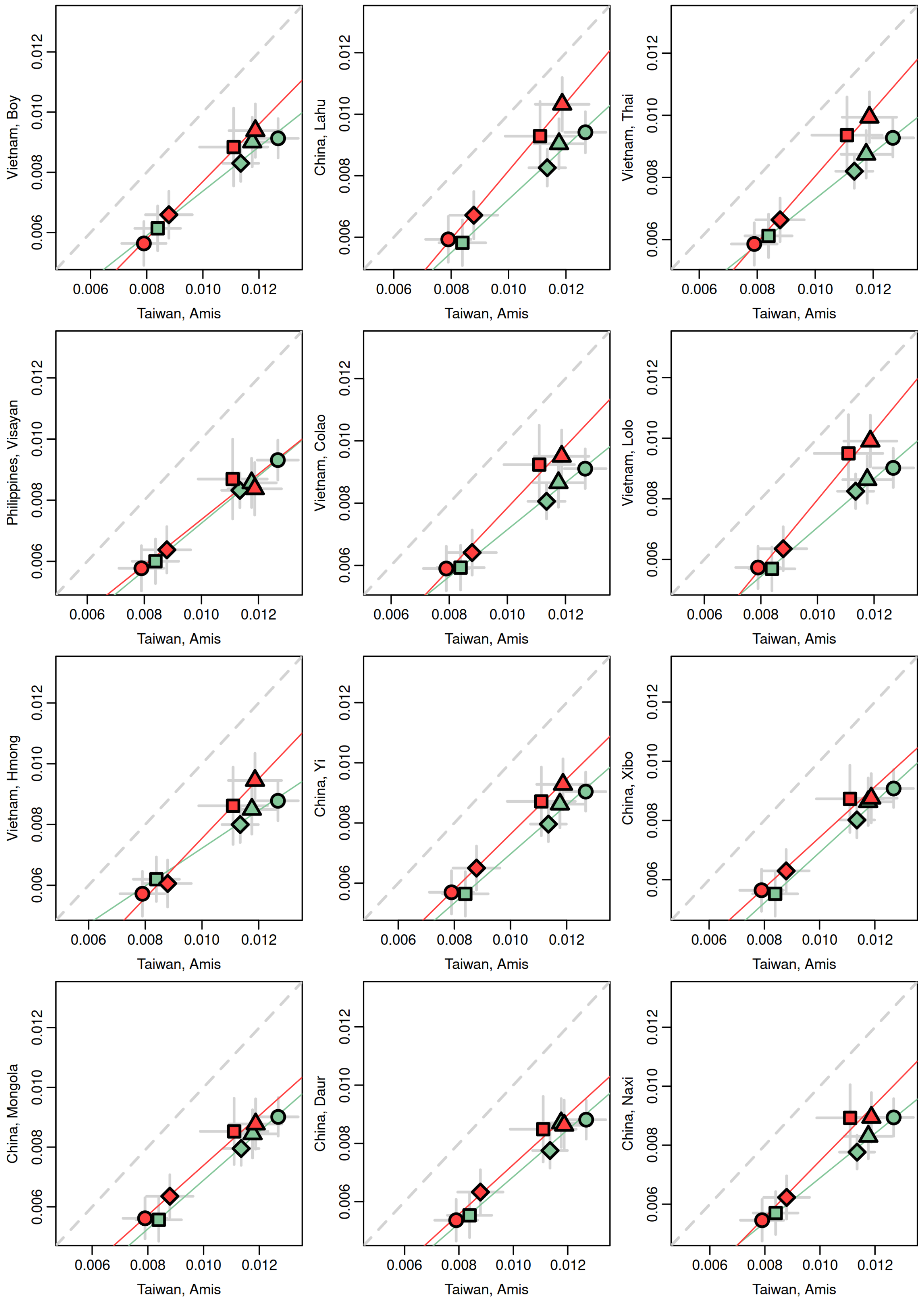
Vietnam, Boy
China, Lahu
Vietnam, Thai
Philippines, Visayan
Vietnam, Colao
Vietnam, Lolo
Vietnam, Hmong
China, Yi
China, Xibo
China, Mongola
China, Daur
China, Naxi
Taiwan, Amis
0.006
0.008
0.010
0.012

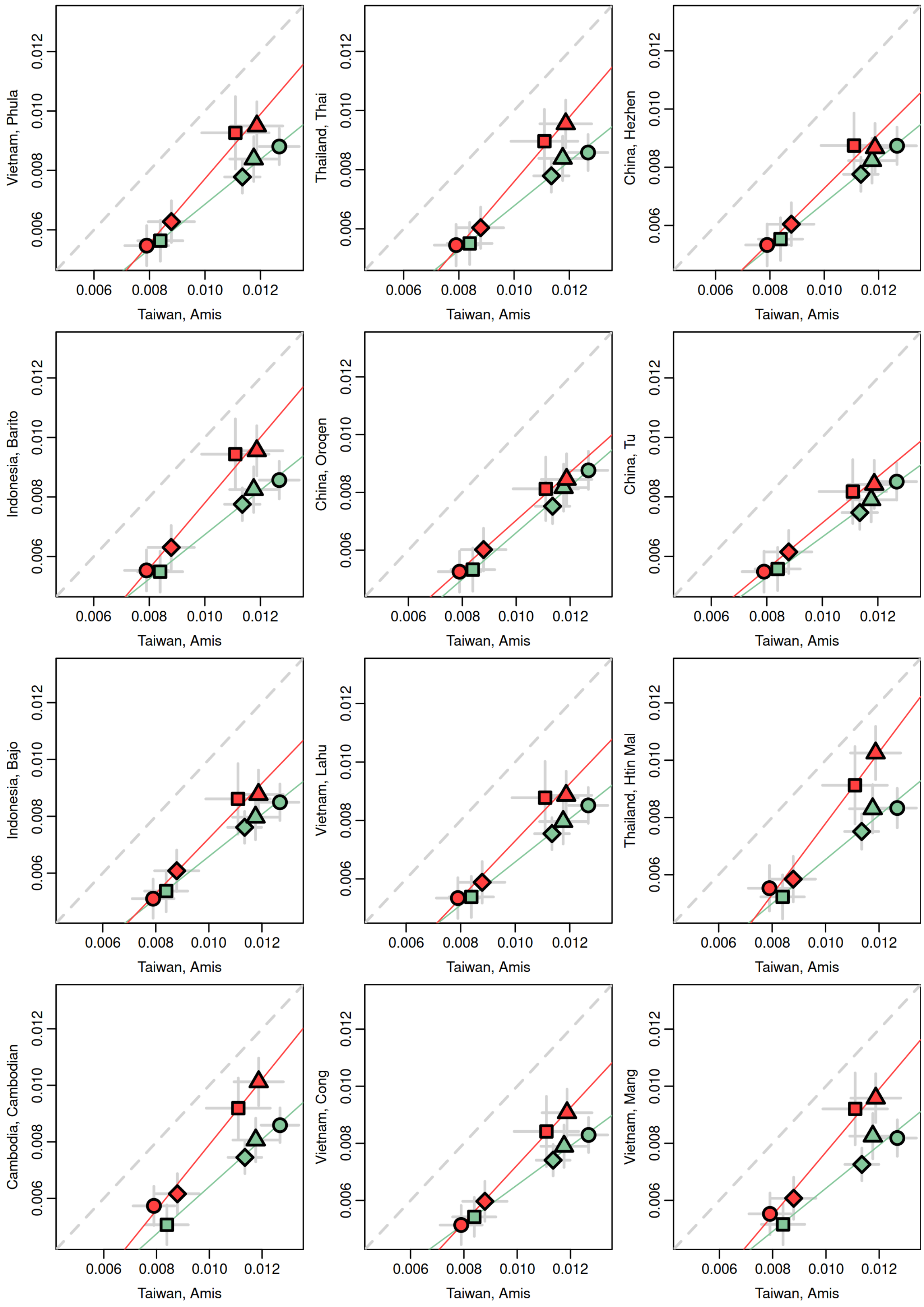

Vietnam, Phula
Thailand, Thai
China, Hezhen
Indonesia, Barito
China, Oroqen
China, Tu
Indonesia, Bajo
Vietnam, Lahu
Thailand, Htin Mal
Cambodia, Cambodian
Vietnam, Cong
Vietnam, Mang
Taiwan, Amis
0.006
0.008
0.010
0.012

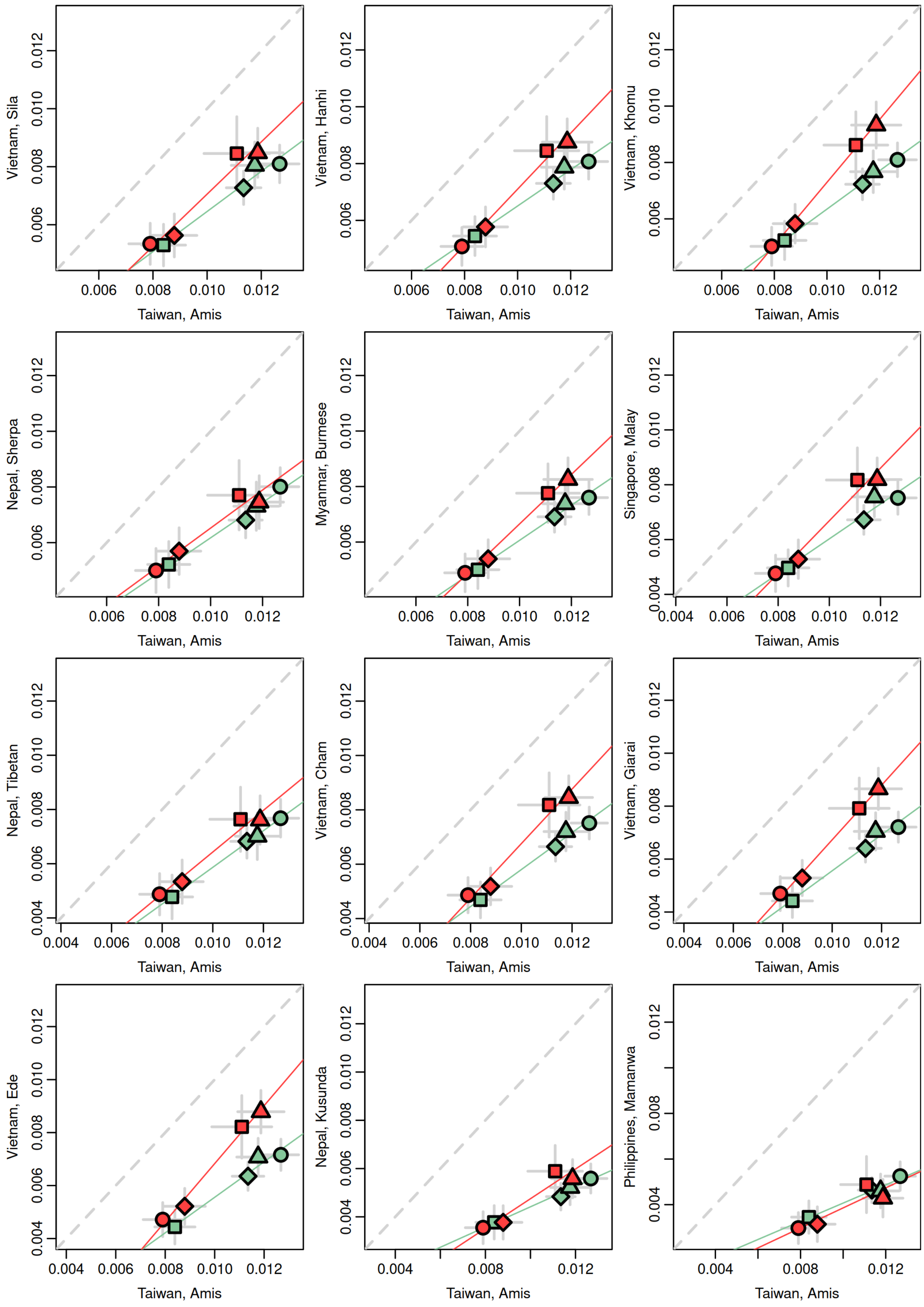

Vietnam, Sila
Taiwan, Amis
Vietnam, Hanhi
Taiwan, Amis
Vietnam, Khomu
Taiwan, Amis
Nepal, Sherpa
Taiwan, Amis
Myanmar, Burmese
Taiwan, Amis
Singapore, Malay
Taiwan, Amis
Nepal, Tibetan
Taiwan, Amis
Vietnam, Cham
Taiwan, Amis
Vietnam, Giarai
Taiwan, Amis
Vietnam, Ede
Taiwan, Amis
Nepal, Kusunda
Taiwan, Amis
Philippines, Mamanwa
Taiwan, Amis
0.004
0.006
0.008
0.010
0.012

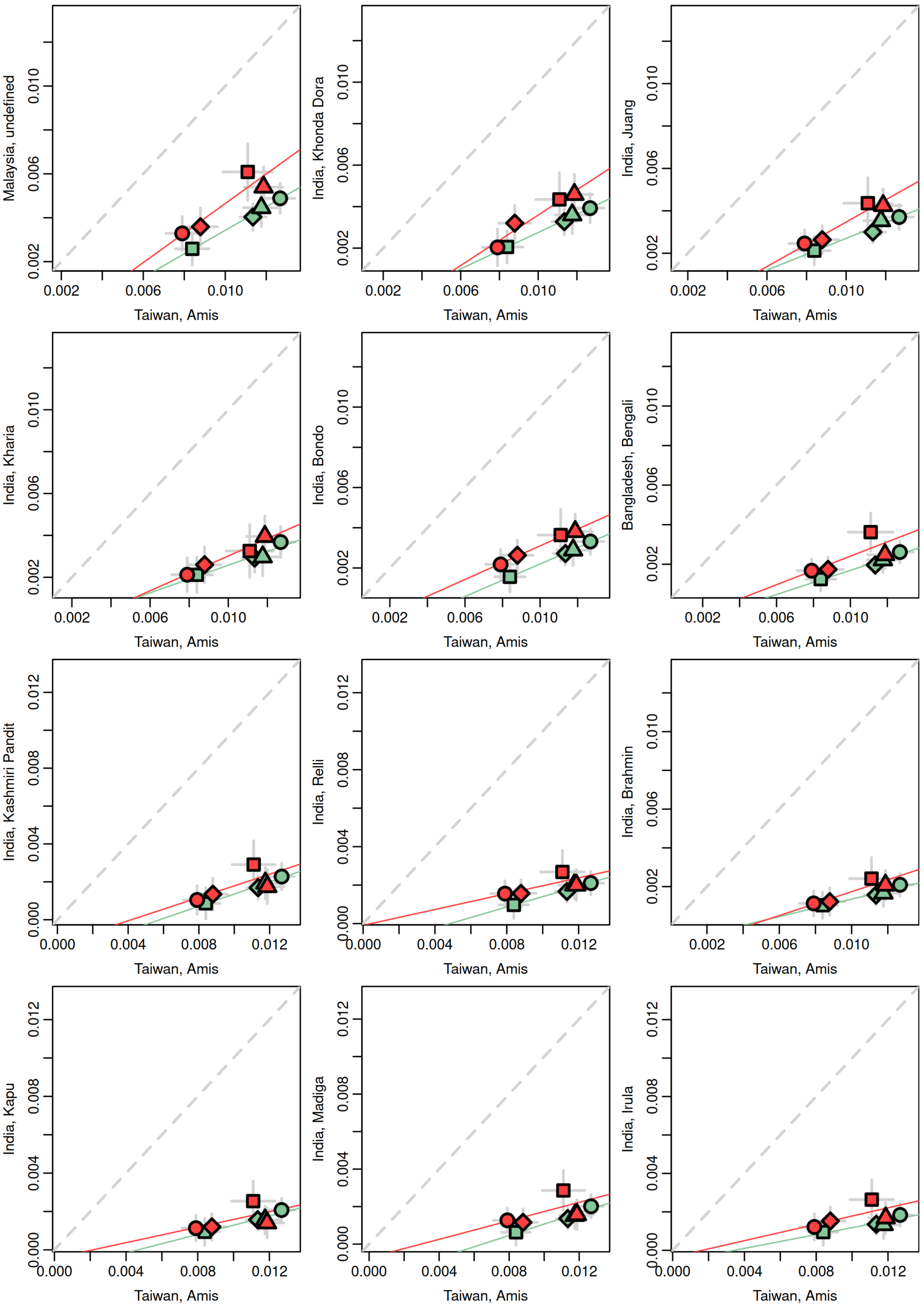
Malaysia, undefined
0.002
0.006
0.010
Taiwan, Amis
India, Khonda Dora
0.002
0.006
0.010
Taiwan, Amis
India, Juang
0.002
0.006
0.010
Taiwan, Amis
India, Kharia
0.002
0.006
0.010
Taiwan, Amis
India, Bondo
0.002
0.006
0.010
Taiwan, Amis
Bangladesh, Bengali
0.002
0.006
0.010
Taiwan, Amis
India, Kashmiri Pandit
0.000
0.004
0.008
0.012
Taiwan, Amis
India, Relli
0.000
0.004
0.008
0.012
Taiwan, Amis
India, Brahmin
0.002
0.006
0.010
Taiwan, Amis
India, Kapu
0.000
0.004
0.008
0.012
Taiwan, Amis
India, Madiga
0.000
0.004
0.008
0.012
Taiwan, Amis
India, Irula
0.000
0.004
0.008
0.012
Taiwan, Amis

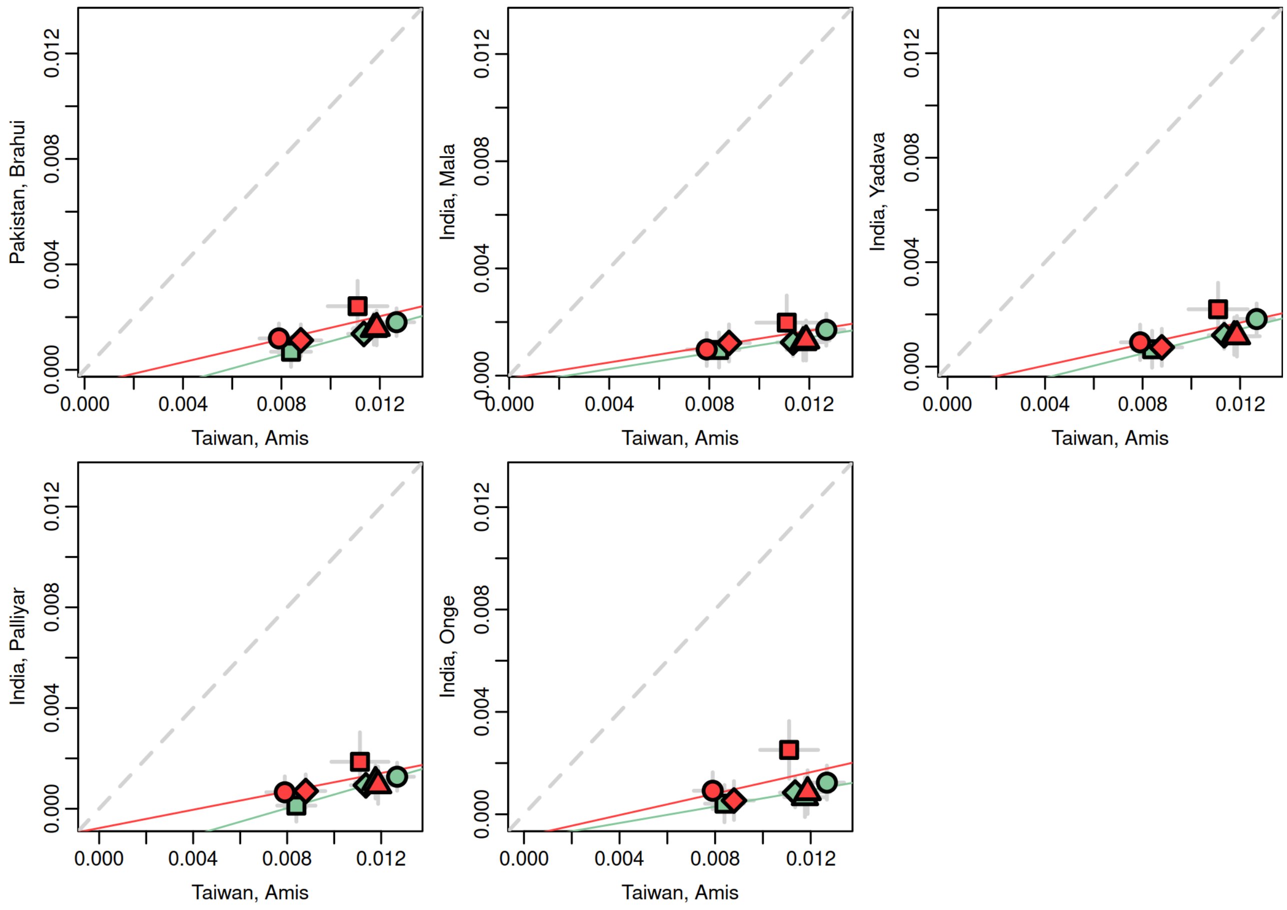

Pakistan, Brahui
India, Mala
India, Yadava
India, Palliyar
India, Onge
Taiwan, Amis
0.000
0.004
0.008
0.012

**Figure S6 - Biplots showing the results of pairs of $f_4$-statistics of the form: *$f_4$(Mbuti, test; New Guinea Highlanders, ancient Wallacea).*** The test groups, shown on the x-y axis label, include ancient groups from mainland Asia, Island southeast Asia, and Oceania that have no discernible Papuan-related ancestry based on the DyStruct analysis. Linear regression lines for the North Moluccas and East Nusa Tenggara individuals are shown in green and red, respectively.

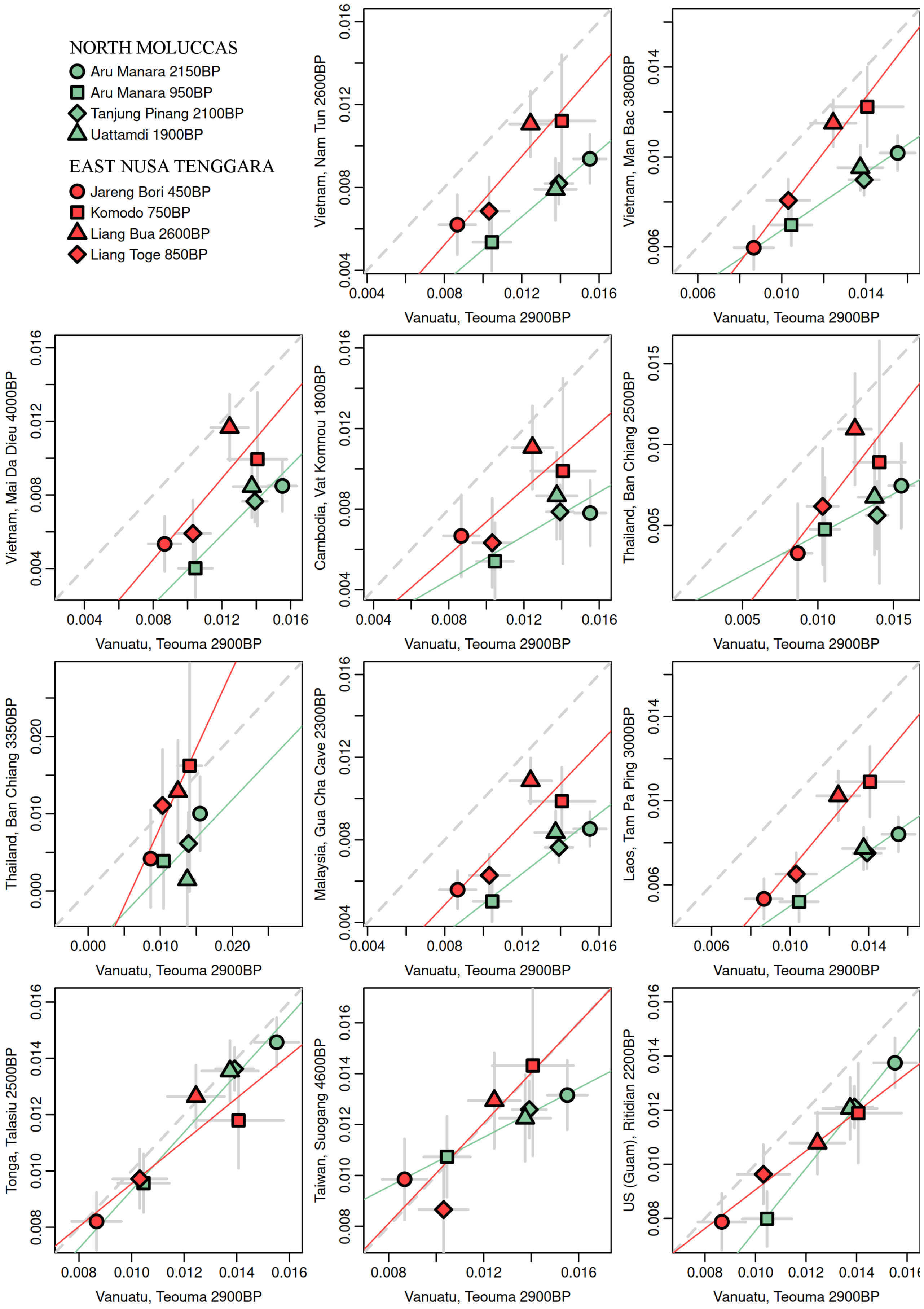

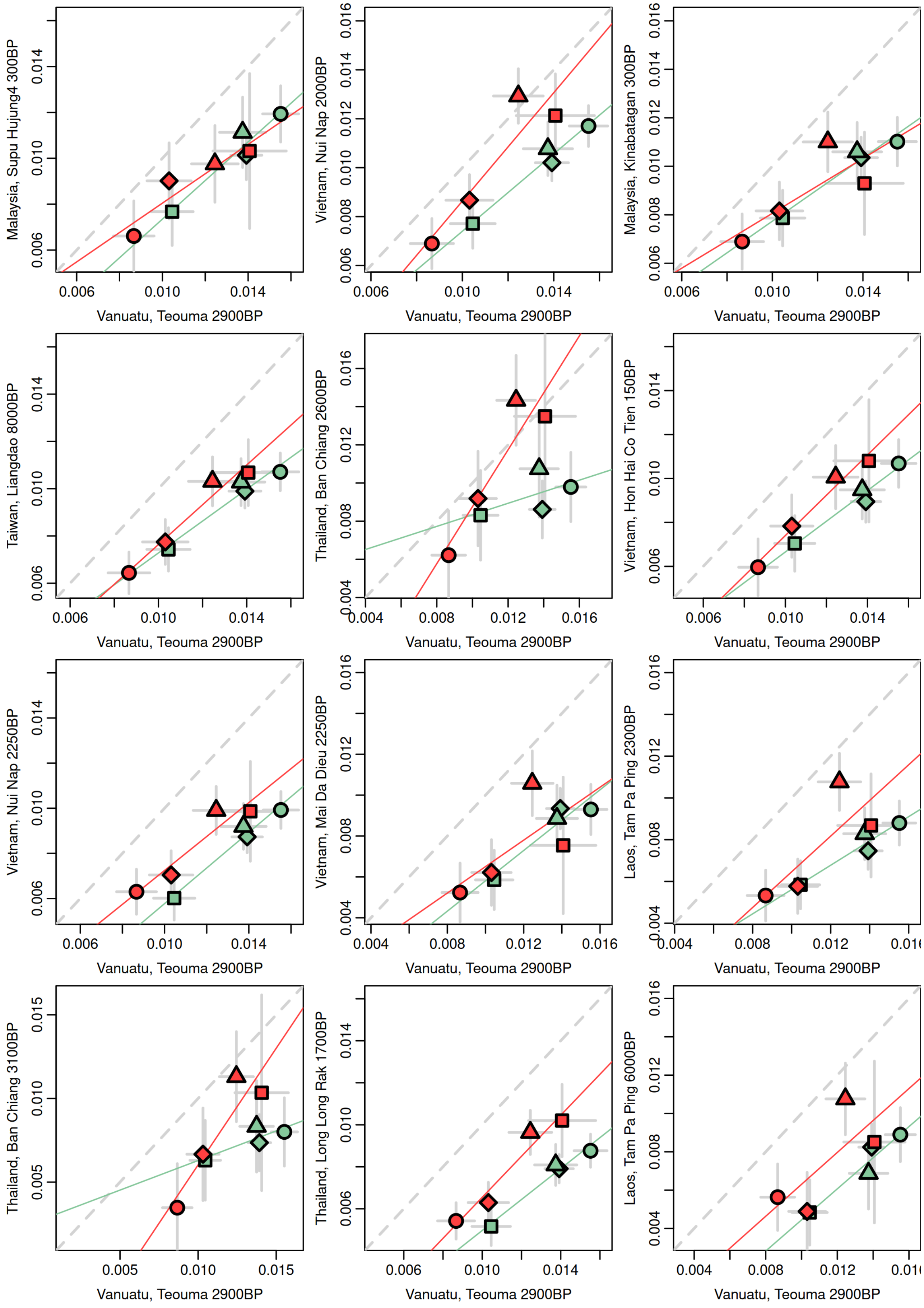
Malaysia, Supu Hujung4 300BP
Vietnam, Nui Nap 2000BP
Malaysia, Kinabatagan 300BP
Taiwan, Liangdao 8000BP
Thailand, Ban Chiang 2600BP
Vietnam, Hon Hai Co Tien 150BP
Vietnam, Nui Nap 2250BP
Vietnam, Mai Da Dieu 2250BP
Laos, Tam Pa Ping 2300BP
Thailand, Ban Chiang 3100BP
Thailand, Long Long Rak 1700BP
Laos, Tam Pa Ping 6000BP
Vanuatu, Teouma 2900BP

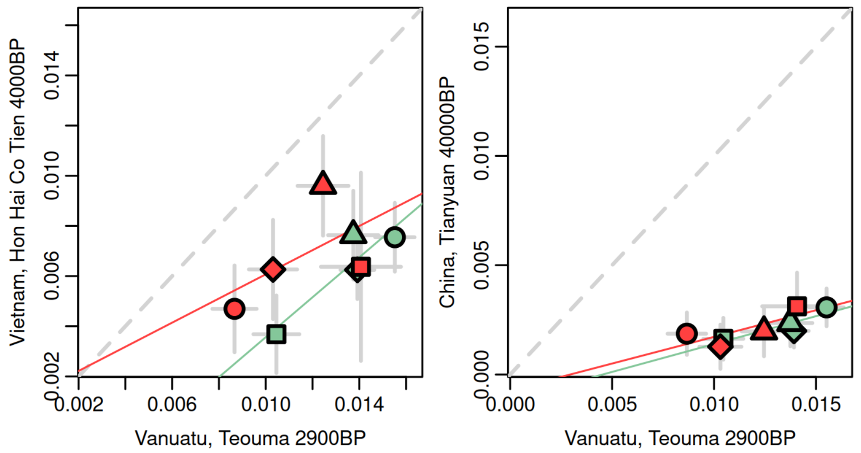

Vietnam, Hon Hai Co Tien 4000BP
0.002
0.006
0.010
0.014
Vanuatu, Teouma 2900BP
China, Tianyuan 40000BP
0.000
0.005
0.010
0.015
Vanuatu, Teouma 2900BP

**Figure S7 - *$f_4$*-statistic of the form *$f_4$(Mbuti, new ancient Wallacean; New Guinea Highlanders, test)* computed for each ancient group from Wallacea separately.** The test groups (shown to the right of each value) consist of Australo-Papuans with no discernable Asian ancestry and a recently published pre-Neolithic individual from Sulawesi (Leang Panninge). Bars indicate two standard errors in both directions. Values in green are not significantly different from zero (|Z| < 2), whereas values in red are significantly different from zero (|Z| > 2). Significant negative results indicate that the new ancient Wallacean shares additional drift with the New Guinea Highlands, while positive results indicate that the new ancient Wallacean shares additional drift with the test population.

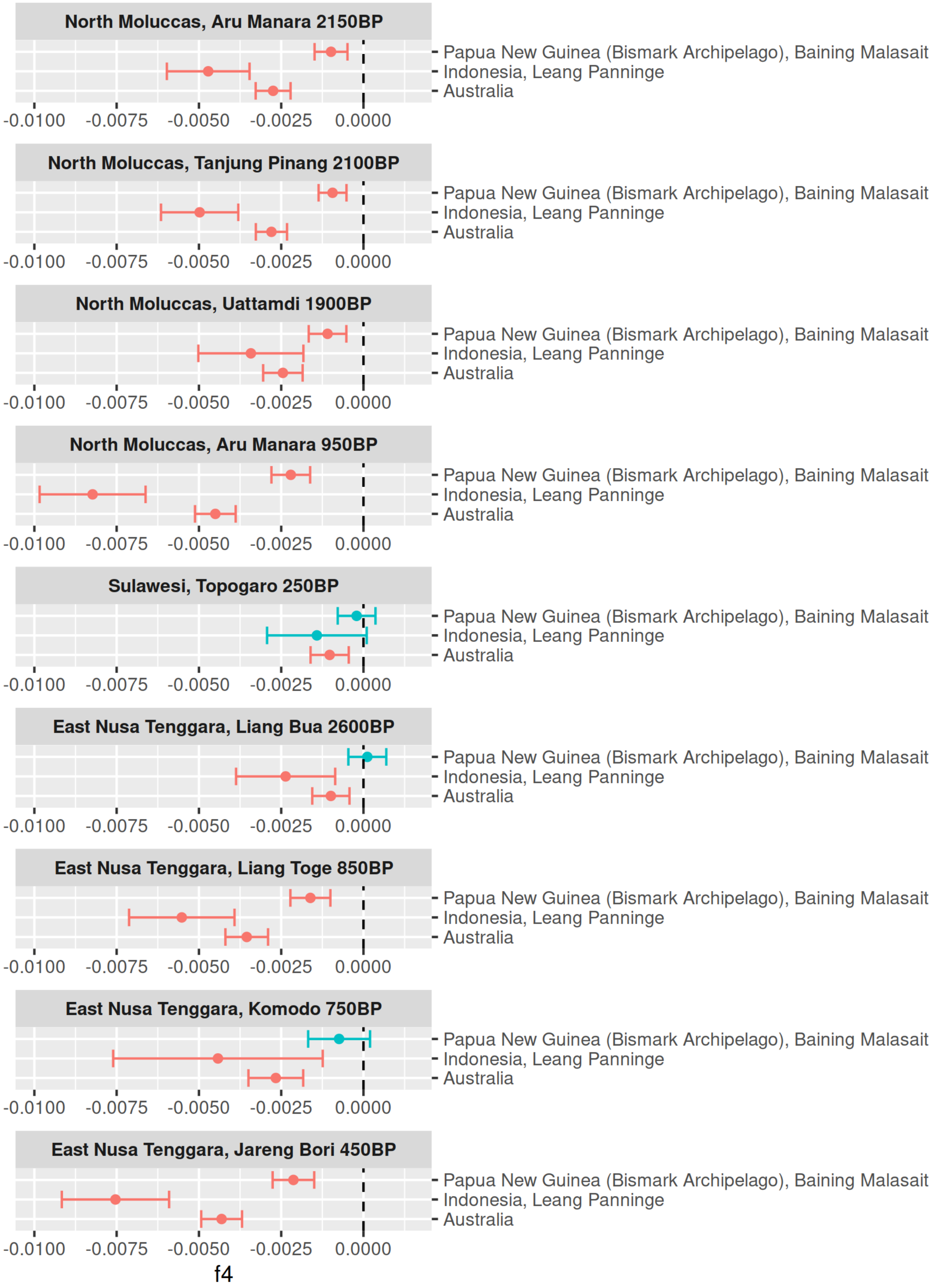

**Figure S8 – Correlation between the amount of Papuan-related ancestry in ancient Wallacean groups and their shared drift with different Australo-Papuan groups, as well as a recently published pre-Neolithic individual from Sulawesi (Leang Panninge).** The Papuan-related ancestry was estimated with qpAdm (see methods) and the values of shared drift correspond to those displayed in Figure S7 (here shown in the y-axis). Grey bars indicate one standard error in each direction.

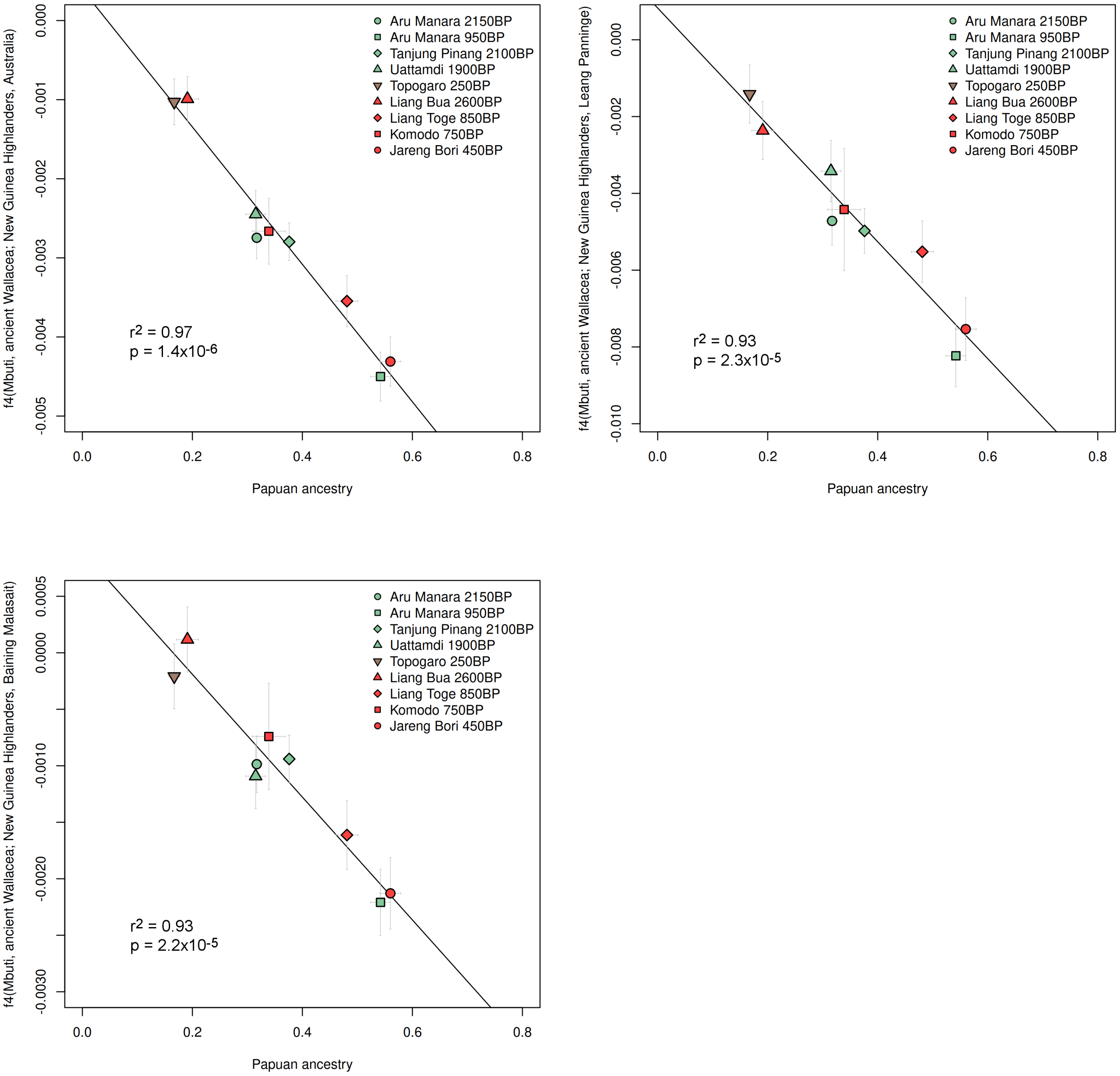

**Figure S9 – Ancestry covariance in ancient Wallaceans.** The number of individuals included in each group is shown in parenthesis, next to the group label. The inferred admixture date (before adding the sample age) and standard errors are shown in generations.

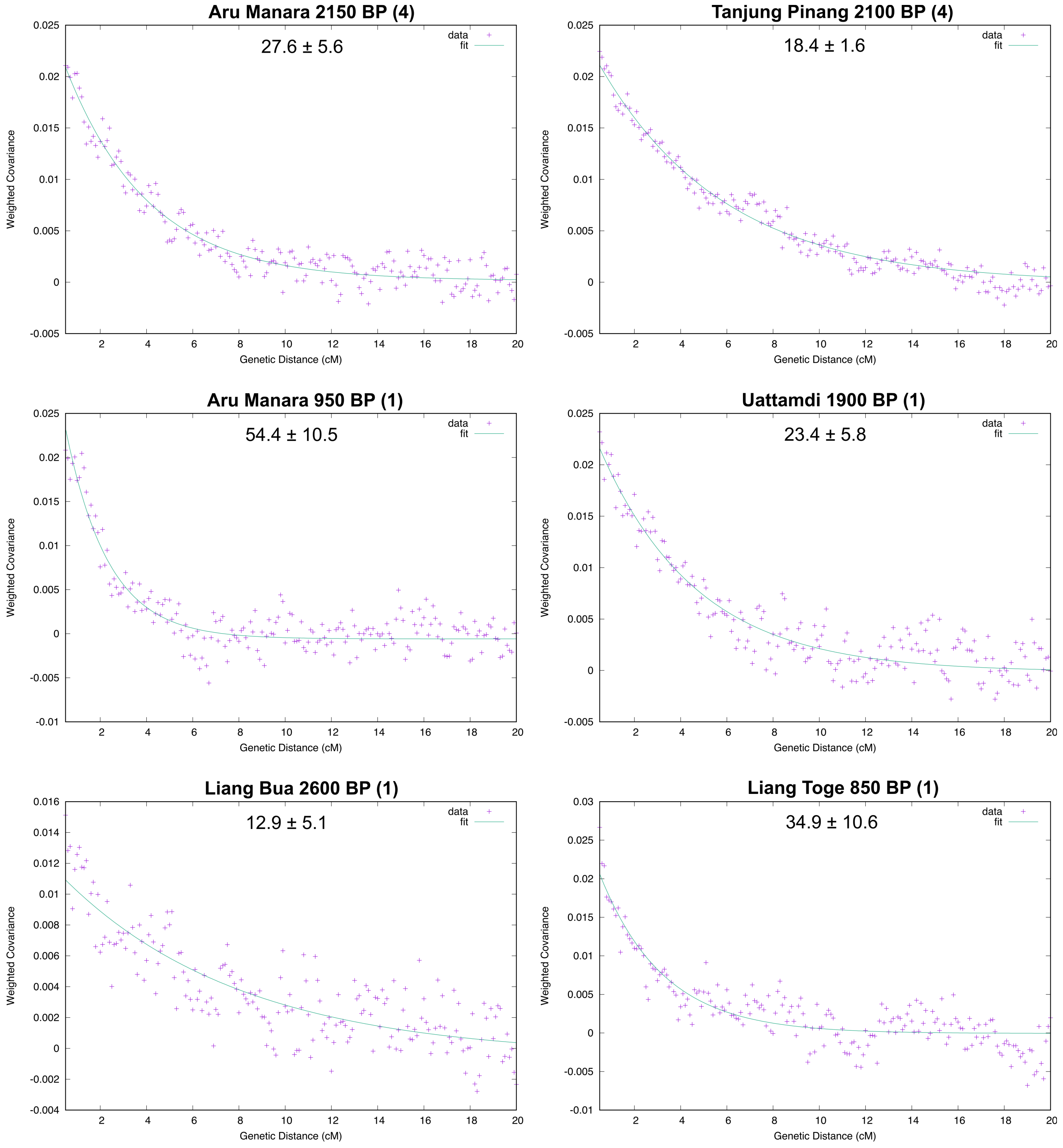

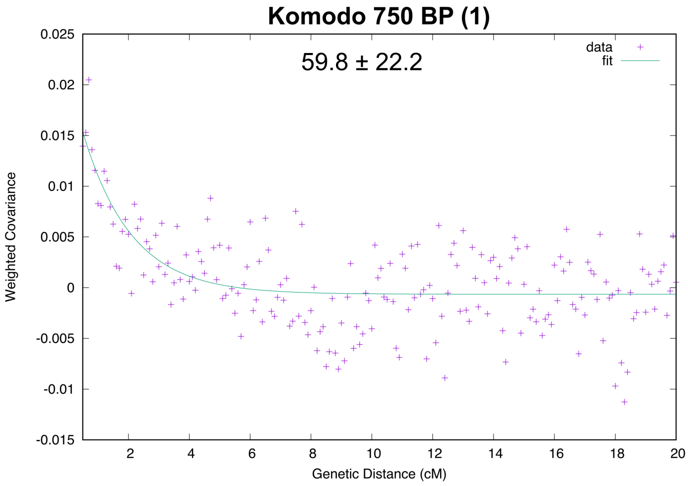
Komodo 750 BP (1)
59.8 ± 22.2
data
fit
Weighted Covariance
Genetic Distance (cM)
0.025
0.02
0.015
0.01
0.005
0
-0.005
-0.01
-0.015
2
4
6
8
10
12
14
16
18
20

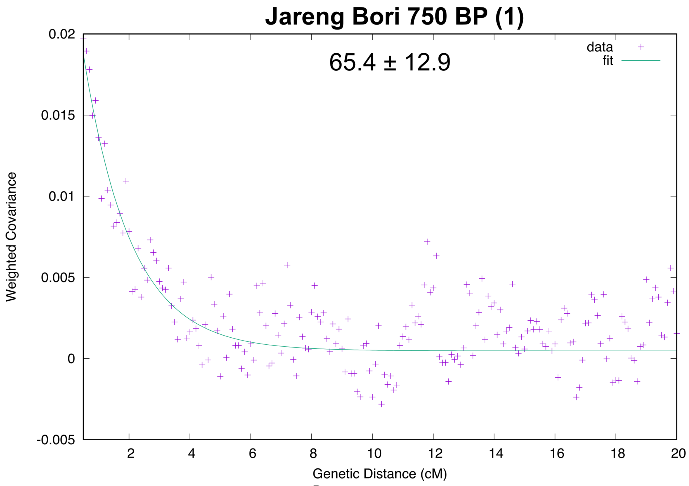
Jareng Bori 750 BP (1)
65.4 ± 12.9
data
fit
Weighted Covariance
Genetic Distance (cM)
0.02
0.015
0.01
0.005
0
-0.005
2
4
6
8
10
12
14
16
18
20

D

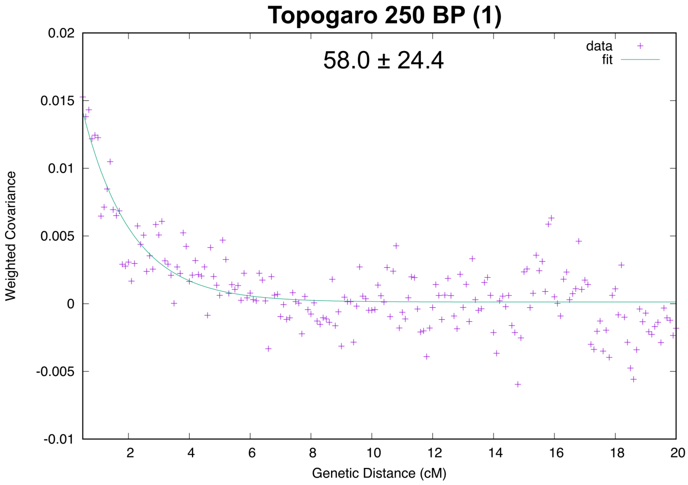
Topogaro 250 BP (1)
58.0 ± 24.4
data
fit
Weighted Covariance
Genetic Distance (cM)
0.02
0.015
0.01
0.005
0
-0.005
-0.01
2
4
6
8
10
12
14
16
18
20

**Figure S10 - Papuan vs Denisova ancestry.** The Papuan-related ancestry was estimated using qpAdm (see methods). The Denisova ancestry is represented by an $f_4$-statistic of the form $f_4$*(Mbuti, Denisova; French, test)*. Grey bars indicate one standard error in each direction.

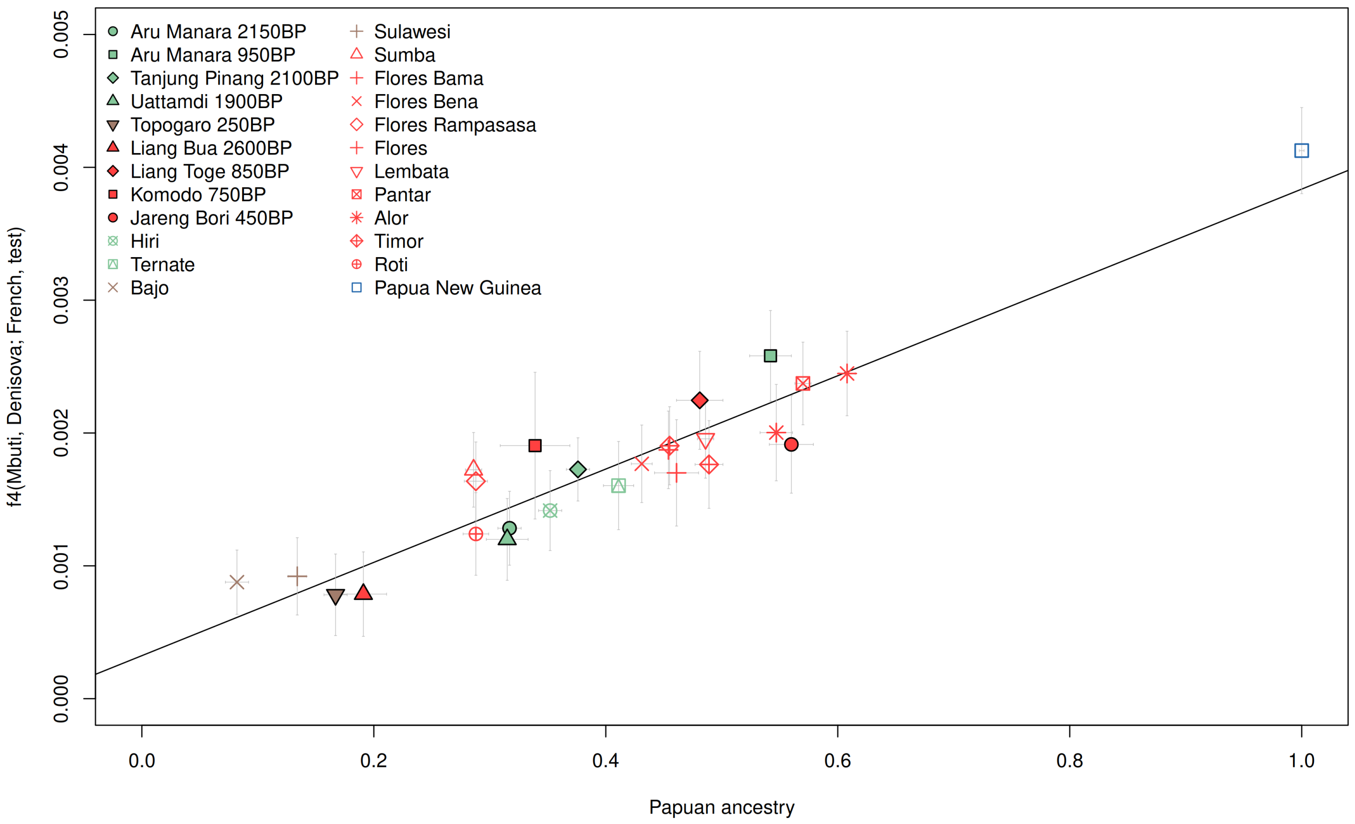
